# CRK2 controls the spatiotemporal distribution of QSK1 at plasma membrane during osmotic stress

**DOI:** 10.64898/2025.12.09.692923

**Authors:** Sunita Jindal, Adam Zeiner, Alexey Bondar, Michaela Neubergerová, Sara Christina Stolze, Anne Harzen, Francisco J. Colina, Sara Liekens, Mirva Pääkkönen, Johannes Merilahti, Ivan Kulich, Roman Pleskot, Hirofumi Nakagami, Michael Wrzaczek

## Abstract

Precise control of intercellular communication is essential for normal growth and stress responses in all multicellular organisms. In Arabidopsis, two membrane-localized receptor like kinases (RLKs), the Cysteine-rich RLK CRK2 and the Leucine-rich repeat (LRR) RLK QSK1 relocalize from the general plasma membrane (PM) to plasmodesmata (PD) in response to osmotic stress. Both these RLKs regulate callose deposition thereby modulating PD permeability. However, unchecked callose deposition can block the PD and disrupt proper intercellular communication. Here, we show that under normal growth conditions, CRK2 phosphorylates and sequesters QSK1 at the general PM, preventing unnecessary callose deposition at PD. We show that osmotic stress-induced enrichment of QSK1 at PD requires functional CRK2 and establish that phosphorylation of QSK1 in its C-terminal region is inhibitory in this process. We propose that osmotic stress triggers dephosphorylation and release of QSK1 from the CRK2-QSK1 complex, enabling its relocalization from general PM to PD, where it promotes stress-induced callose deposition. Subsequently, CRK2 relocalizes to PD where it negatively influences callose deposition. Our work reveals a tightly coordinated distribution of QSK1 and CRK2 at PM, establishing a dynamic gating mechanism that balances growth and stress responsiveness.

## Introduction

In natural habitats, plant water uptake is compromised by salinity, drought and extreme temperatures leading to osmotic stress which limits normal growth and development. On the physiological level, osmotic adjustments are made by closing stomata, growth inhibition, hydraulic modifications and changes in root system architecture. On the molecular level, rapid signaling events are triggered at the plasma membrane (PM) and include activation of phosphorylation networks, activation of NADPH oxidases (respiratory burst oxidase homologs, RBOHs) and ROS burst^1^, Ca^2+^ influx^2–4^ and changes in lipid signaling, for example, accumulation of phosphatidic acid (PA)^5–8^. Another key regulatory element of osmotic stress signaling is the deposition of callose^9–11^. Callose, a β-1,3-glucan polysaccharide is deposited primarily in the neck region of plasmodesmata (PD) and controls symplastic connectivity through gating the PD aperture^12,13^. Symplastic connection regulates cell division, expansion and organ formation^14,15^ which could provide growth adaptation especially during extreme stress conditions. For example, mutants accumulating higher levels of callose have reduced primary root (PR) length^9,16^ and altered lateral root (LR) patterning^12,17^. Therefore, adaptations in the root phenotype during osmotic stress can at least partly result from the changes in symplastic connection.

The amount of callose at PD changes dynamically throughout plant growth, development and during stress conditions. The main determinants of callose amounts are callose synthases (CALSs) and β-1,3-glucanases (PdBGs), responsible for callose synthesis and degradation, respectively^18^. Several PD-localized proteins (PDLPs), PD-localized chaperons such as PD-callose binding proteins (PDCBs) participate in maintaining callose homeostasis by recruiting or stabilizing CALSs or PdBGs^18–20^. In recent years, receptor-like kinases (RLKs) have emerged as regulating callose accumulation during abiotic stress^9,21,22^. We previously showed that CRK2, a cysteine-rich RLK (CRK) is involved in the regulation of callose deposition and positively regulates seed germination during salinity stress^9^. The mutant *crk2* exhibits an overall small phenotype and deposits higher callose. CRK2 relocalizes to PD in response to salt and mannitol treatments and also assembles in nanodomain-like clusters upon flg22 and H_2_O_2_ exposure^9^. Concurrently,^21^ showed that Qian Shou kinase 1 (QSK1), a leucine-rich repeat RLK (LRR-RLK) is recruited to PD upon osmotic stress and the *qsk1* mutant accumulates lower callose levels and shows altered root response to mannitol.

PM components undergo dynamic re-organization upon perception of various stimuli and cluster into nanodomains^23^ to facilitate the assembly of signaling complexes and providing signal specificity^24,25^. Membrane nanodomains are hallmarked by specific lipid and protein compositions. Scaffold proteins such as Remorins and Stomatin/Prohibitin/Flotillin/HflK/C (SPFH) domain-containing protein bridge lipids and membrane receptors, whereas RLKs are dynamically positioned in nanodomains^26,27^. Cytoskeletal elements and cell wall components further regulate nanodomain mobility and spatial distribution^27^. The re-organization of membrane proteins is often coupled to a phosphorylation switch. For example, in resting cells, BRI1 (Brassinosteroid Insensitive 1)-Associated receptor Kinase 1 (BAK1), an LRR-RLK, is evenly distributed across the PM, with its catalytic pocket shielded by BAK1-Interacting Receptor-like kinases (BIRs)^28^. Ligand perception activates the BRI1-BAK1 or FLAGELLIN-SENSITIVE 2 (FLS2)-BAK1 signaling axes, triggering growth or immune signaling, respectively. This involves a phosphorylation-dependent shift of BAK1 from resting to active state and the formation of receptor nanodomains^29^. Likewise, the distribution of Remorin^30,31^ and QSK1^21^ across the PM is dependent on their phosphorylation status.

Similar to BAK1 as a shared co-receptor in growth and immune signaling, QSK1 forms several protein complexes. For instance, the SIRK1-QSK1 module regulates aquaporins where QSK1 acts as a co-receptor of the RLK Sucrose-Induced Receptor Kinase 1 (SIRK1)^32,33^. QSK1 also associates with pattern recognition receptor (PRR)-RBOHD complexes involving FLS2 and Elongation Factor Tu Receptor (EFR) RLKs^34^. QSK1 also regulates the vacuolar Two-pore potassium channel 1 (TPK1) during stomatal closure by phosphorylating TPK1 with involvement of an unidentified receptor kinase^35^. Similarly, QSK1 associates with the transceptor nitrate transporter NRT1.1 in regulating PM Arabidopsis H^+^-ATPase 2 (AHA2)^36^.

While the re-organization of individual PM components is extensively studied, how different RLKs coordinate their spatiotemporal behavior and how the shared components are exchanged between the complexes is much less understood. CRK2 and QSK1 are both primarily present at the general PM under normal growth conditions but both relocalize to PD upon osmotic stress. We previously identified that QSK1 associates with CRK2^9^. However, the functional link between these two RLKs in osmotic stress response was not explored. Here, we elucidate their connection in fine-tuning callose deposition and plant responses to osmotic stress. We show that CRK2 controls the plasmodesmal relocalization of QSK1 and both RLKs antagonistically regulate callose deposition during osmotic stress responses. Our work highlights that the dynamics of the re-organization of QSK1 and CRK2 in response to osmotic stress is spatiotemporally coordinated and involves a phosphorylation switch.

## Results

### Role of CRK2 and QSK1 in plant growth and osmotic stress response

The growth phenotype and osmotic stress responses of the *crk2* and *qsk1* have been previously described^9,21,37^. We generated the *crk2 qsk1* double mutant by crossing *crk2* and *qsk1* single mutants and characterized the growth phenotype under normal growth conditions salinity stress. As reported previously^9^, *crk2* displayed overall smaller rosette, shorter primary root (PR) and lower lateral root (LR) density (Fig. 1A-C, Table S1). The *qsk1* rosette area was comparable to Col-0 wild type (WT), while in *crk2 qsk1*, the small rosette phenotype of *crk2* was rescued with a slight increase in comparison to the WT (Fig. 1A, Fig. S1A). While *crk2* exhibited the shortest PR, *qsk1* and *crk2 qsk1* also displayed shorter PR compared to the WT (Fig. 1B, Fig. S1B). Similarly, LR density was lower in all the tested mutants compared to WT (Fig. 1C). Overall, the observable phenotypic traits of *crk2 qsk1* were more similar to either WT or *qsk1* rather than *crk2*. The partial reversion of *crk2* phenotypes in *crk2 qsk1* mutant was also observed at the level of stress responses. We tested WT, *crk2*, *qsk1* and *crk2 qsk1* seeds for salt-sensitivity during seed germination and observed that, *crk2* seeds showed hypersensitivity to salt as reported previously^9^, whereas *qsk1* and *crk2 qsk1* seed germination was comparable to WT (Fig. 1D). Taken together, these results suggest, that CRK2 and QSK1 functionally interact to regulate plant growth and stress responses, and moderation of *crk2* phenotype by introduction of additional *qsk1* mutation points towards antagonistic roles for those two RLKs.

**Figure 1.**
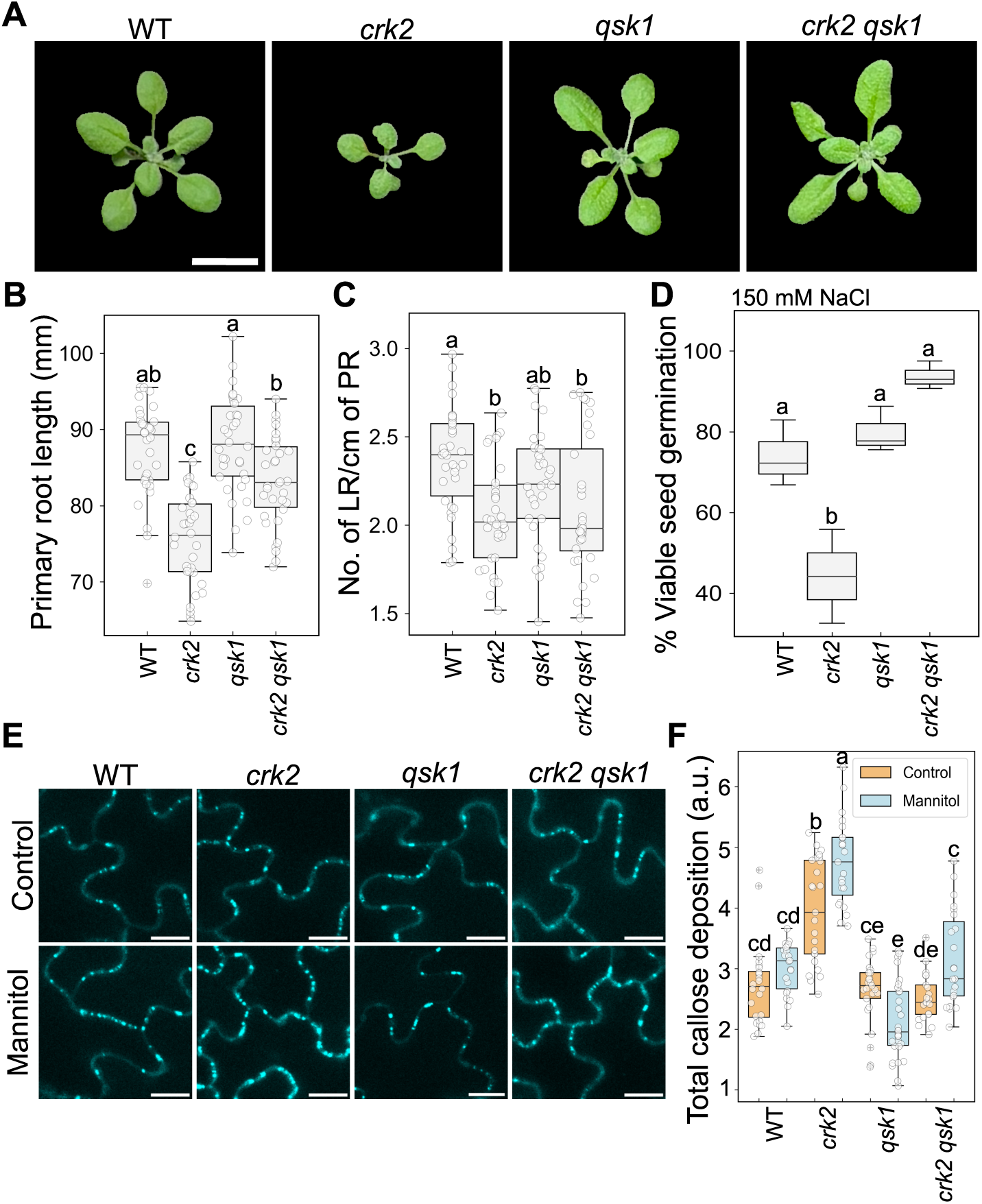
CRK2 and QSK1 regulate plant growth and play antagonistic roles in osmotic stress-induced callose deposition. (A) Rosette of 2-week-old Arabidopsis plants grown in long-day conditions. Mutation of *QSK1* reverts small growth phenotype of *crk2* in the double mutant *crk2 qsk1*. Scale bar 1 cm. (B) Primary root (PR) length of 12-days-old plants. (C) Lateral root (LR) number per cm of PR (LR density). (D) Percentage of viable seed germination on media containing NaCl. The hypersensitive response of *crk2* mutant on salt-containing media was reverted in the *crk2 qsk1* double mutant. The germination efficiency was normalized according to the respective control plates. (E-F) Visualization and quantification of callose deposition in aniline blue-stained leaf epidermal cells of Arabidopsis. Roots of 2-week-old plants were subjected to 50 mM mannitol for 7 hours before imaging in the treated groups. Images acquired with a laser-scanning confocal microscope under the DAPI filter with emission wavelength 461 nm and laser wavelength 405 nm. Detection wavelength 430-530 nm; laser transmissivity 5.0 %; objective lens 60.0 X oil immersion, scale bar 10 µm. The boxes represent the interquartile range (25th–75th percentile), the horizontal line indicates the median, the whiskers extend to 1.5× the IQR, and dots represent individual data points. Panel B, n=34, Kuskal-Wallis test followed by Dunn’s multiple comparison; Panel C, n=34, one-way ANOVA, Tukey’s test; Panel D, n=3, one-way ANOVA, Tukey’s test; Panel E, n≥22, two-way ANOVA, Tukey’s multiple comparison test. Alphabets represent statistically significant differences. See also Fig. S1, Table S1 and Table S2.

### CRK2 and QSK1 fine-tune callose deposition in response to osmotic stress

The mutants *crk2* and *qsk1* have been previously reported to accumulate altered levels of callose^9,21^. We quantified callose accumulation in WT, *crk2*, *qsk1* and *crk2 qsk1* seedlings by aniline blue staining followed by microscopy-based detection of the aniline blue fluorescence signal (Fig. 1E-F, Fig. SC-D, Table S2)^9,38,39^. Since osmotic stress is a common component of various abiotic stresses, including salinity, drought, waterlogging, and extreme temperatures, we used mannitol to specifically impose osmotic stress without introducing additional ionic effects. Arabidopsis plants were treated with mannitol for 7 hours before aniline blue staining and imaging. We quantified total callose deposition (Fig. 1F) by taking into consideration mean fluorescence intensity of the spots and the number of deposits (Fig. S1C-D, Table S2). In the *crk2* mutant, total callose deposition was higher than the WT in control conditions which further increased upon mannitol treatment (Fig. 1F). This suggested a negative influence of CRK2 on callose accumulation during steady-state as well as osmotic stress conditions. In the *qsk1* mutant under control conditions, total callose was comparable to the WT (Fig. 1F). Surprisingly, mannitol-treated *qsk1* showed a lower total callose signal compared to normal growth conditions suggesting that QSK1 might not play a role in basal callose deposition but positively regulates osmotic stress-induced callose deposition (Fig. 1F). Interestingly, the reciprocal impact of CRK2 and QSK1 seemed to negate each other in the *crk2 qsk1* mutant as this mutant accumulated similar callose levels as WT under control and mannitol treatment (Fig. 1F, Fig. SC-D). Together, these findings suggest opposing roles of CRK2 and QSK1 in regulating osmotic stress-induced callose deposition.

### Functional CRK2 is essential for osmotic stress-induced enrichment of QSK1 at PD

CRK2 and QSK1 were shown to relocalize to PD in response to osmotic stress^9,21^ and QSK1 was identified as an *in planta* interactor of CRK2^9^. We hypothesized that their interaction might be necessary for the enrichment of both proteins at PD. Therefore, we tested whether the absence of functional CRK2 or QSK1 would impact the recruitment of the respective other RLK to PD. We generated transgenic plants expressing QSK1 fused to mCherry under native *QSK1* promoter in *crk2* and WT backgrounds and imaged leaf epidermal cells. Interestingly, while QSK1 enriched at PD following mannitol or NaCl treatment in WT background, this enrichment was abolished in *crk2* background (Fig. 2A-B, Fig. S2A, Table S3). In QSK1-mCherry/WT, the enrichment index, which is the fluorescence intensity ratio between the enriched spot (PD) and an adjacent non-enriched spot on the bulk PM, increased from 0.940 to 1.693 upon mannitol treatment. It did not increase for QSK1-mCherry/*crk2* and remained approximately 1.0 for both control and mannitol treatment (Fig. 2B). Since callose is particularly present at PD, it is used to mark PD by aniline blue staining^9,38,39^. To test if the enriched QSK1-mCherry spots following osmotic stress treatment corresponded with PD, we simultaneously applied osmotic stress and aniline blue staining. The co-localization of aniline blue-stained callose with enriched mCherry signal supported that the punctate pattern of QSK1-mCherry correspond to PD (Fig. S2B). Consistently, the genetic cross between QSK1-mCherry/*crk2* and CRK2-mVenus/*crk2* rescued QSK1 enrichment at PD in response to mannitol treatment (Fig. 2C). At this timepoint after mannitol treatment, we did not observe PD enrichment for CRK2 possibly due to a lag in the PD relocalization of CRK2 (Fig. 3B). The mutation D450N in the catalytic core of CRK2 kinase domain (CRK2^D450N^) abolishes the phosphorylation activity of CRK2, and CRK2^D450N^ is not able to relocalize to PD^9^. To determine whether the kinase activity of CRK2 was also essential for the recruitment of QSK1 at PD, we crossed QSK1-mCherry/*crk2* with CRK2^D450N^-mVenus/*crk2* plants. In the resulting plants, QSK1 failed to relocalize to PD in response to mannitol treatment (Fig. 2D). To investigate whether CRK2 requires QSK1 for its own PD localization, we stably expressed CRK2-mVenus under native *CRK2* promoter in WT and *qsk1* backgrounds. Interestingly, we found that CRK2 relocalized to PD upon mannitol treatment without functional QSK1 (Fig. 2E, F). These results established that enzymatically active CRK2 is essential for the relocalization of QSK1 to PD in response to osmotic stress. However, the relocalization of CRK2 is independent of QSK1 pointing to an upstream role of CRK2 in PD-targeting of itself and QSK1.

**Figure 2.**
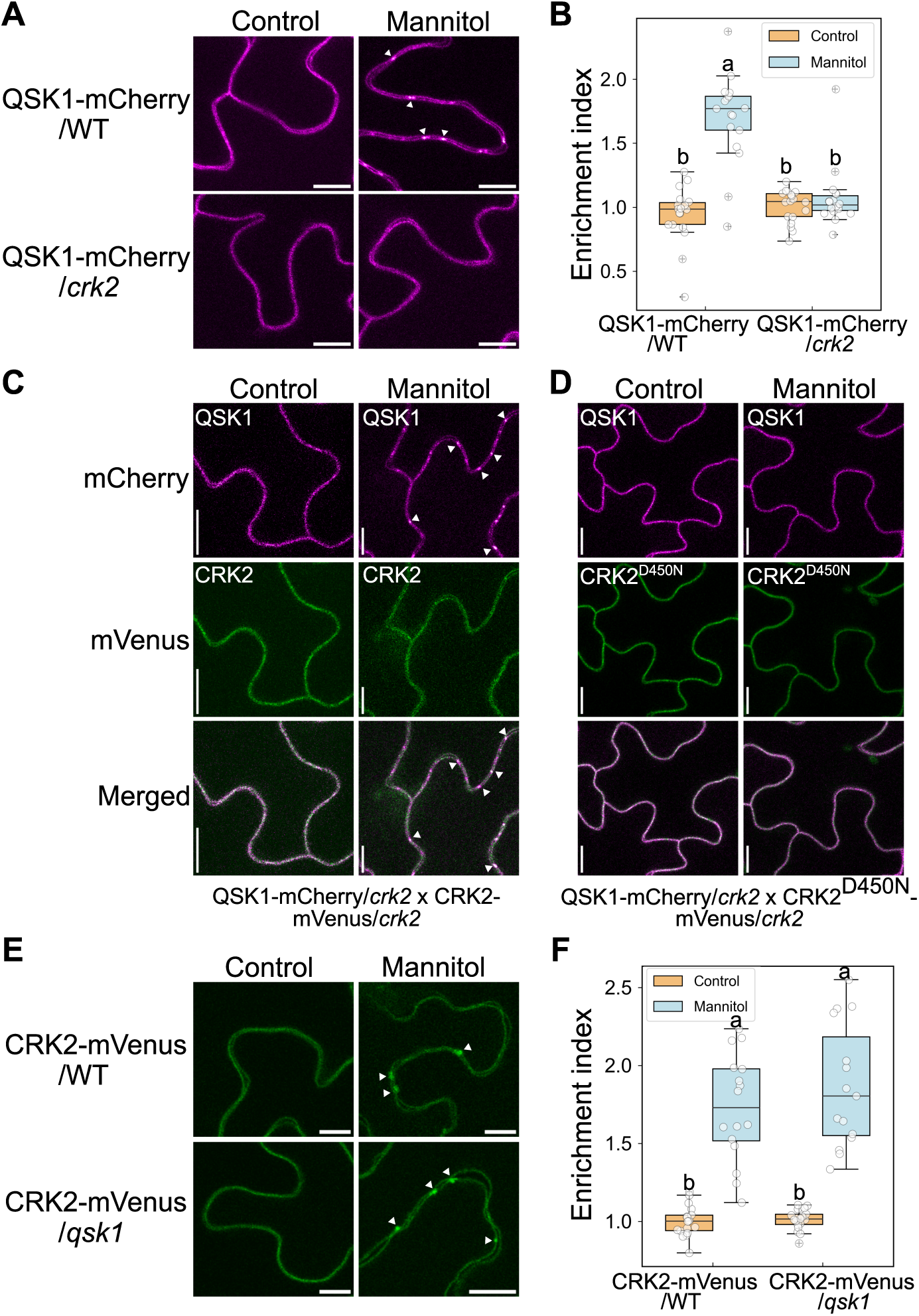
Functional CRK2 is required for osmotic stress-induced enrichment of QSK1 at plasmodesmata. (A) Relocalization QSK1-mCherry in Arabidopsis leaf epidermal cells. Detached leaves from plants expressing QSK1-mCherry in WT or *crk2* mutant backgrounds were treated with 0.4 M mannitol. The abaxial surface was visualized and imaged with a laser-scanning confocal microscope after 30 min of treatment. QSK1 failed to enrich in the *crk2* mutant background. (B) Boxplot showing the extent of PD enrichment of QSK1 which is signal intensity ratio between PD and PM ROI. (C) The disruption of QSK1 enrichment in the *crk2* mutant background was rescued by complementing QSK1-mCherry/*crk2* plants with CRK2-mVenus (QSK1-mCherry/*crk2* crossed with CRK2-mVenus/*crk2*). (D) The kinase inactive CRK2^D450N^ failed to complement this function in the cross of QSK1-mCherry/*crk2* and CRK2^D450N^-mVenus/*crk2* plants. (E) The mutation in *QSK1* did not impact the relocalization of CRK2 in the CRK2-mVenus/*qsk1* plants. Images are acquired after 60 min of 0.4 M mannitol treatment. (F) Boxplot showing enrichment index related to panel E. The images were captured with 60.0 X oil immersion lens. Laser and emission wavelengths were 561 nm and 610 nm, respectively for mCherry; 514 nm and 527 nm for mVenus. Detection wavelengths were 570-620 nm and 530-630 nm for mCherry and mVenus, respectively. Laser transmissivity in the shown images is 7.0%-15% which was kept constant between the comparisons. Scanning of signals were done in a sequential manner to avoid signal cross-contamination between the channels. Scale bar 10 µm. At least 3 independent transgenic lines were tested for each genotype. Boxplots as described in Figure 1. Panels B and F, n ≥ 15, two-way ANOVA, Tukey’s test. See also Fig. S2 and Table S3.

**Figure 3.**
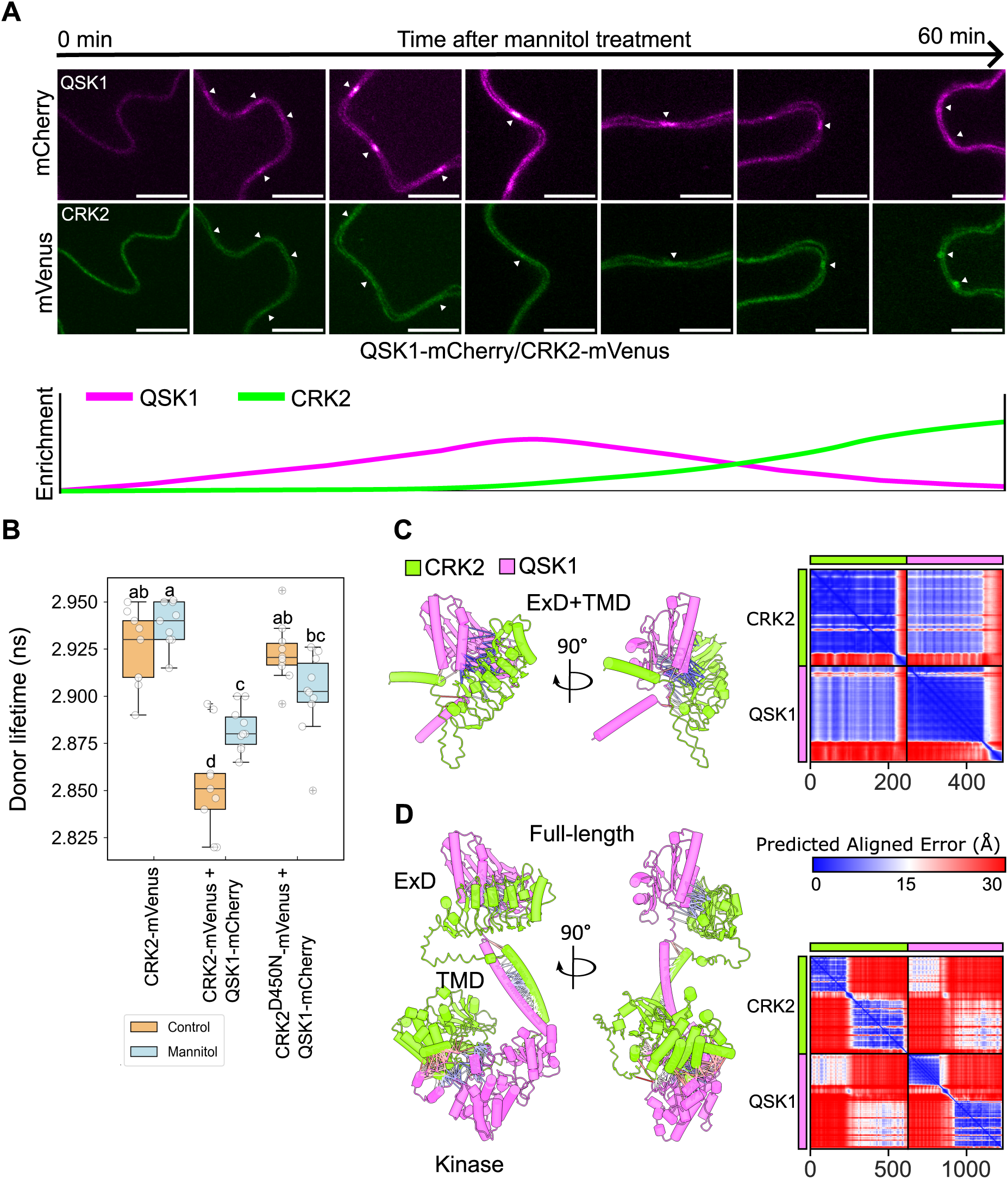
QSK1 and CRK2 are present at the general plasma membrane in a pre-formed complex and relocalize and enrich at plasmodesmata in a sequential manner during osmotic stress. (A) Arabidopsis leaf epidermal cells showing relocalization of QSK1 and CRK2. The images shown are captured within a window of 0-60 min. QSK1 enrichment takes place first and reaches maximum enrichment at approximately 30 min of treatment and subsequently starts to decline. Objective lens 40.0 X oil immersion. Treatment and laser settings as described for Figure 2. Scale bar 10 µm. (B) FRET-FLIM analysis of CRK2-mVenus (donor-only), CRK2-mVenus+QSK1-mCherry (donor-acceptor) and CRK2^D450N^-mVenus+QSK1-mCherry (donor-acceptor negative control) plants with or without mannitol treatment. A lower mVenus lifetime in the donor-acceptor pair CRK2-mVenus+QSK1-mCherry indicated the presence of a preformed complex of CRK2 and QSK1. An increase in the mVenus lifetime in this pair after mannitol treatment suggested dissociation of the complex. Boxplot as described in Figure 1. n ≥ 9. Two-way ANOVA, Tukey’s test. (C- D) AlphaFold2 prediction of interaction between CRK2 and QSK1 extracellular and transmembrane domains (ExD+TMD) (C) and full-length proteins (D). Contacts under 6 Å between the two proteins are shown. Lines are colored according to the PAE values, with most contacts showing high confidence. See also Fig. S3, Table S4 and Table S5.

### QSK1 and CRK2 enrich at PD in a sequential manner following osmotic stress

RLKs are dynamically re-organized at the PM in response to various internal and external cues^24,25^. Despite the dependence of QSK1 on CRK2 in its relocalization to PD, we observed a temporal disconnection in their relocalization. QSK1 accumulation at PD peaked within 30-45 min of salt or mannitol treatment and subsequently the punctate pattern faded with almost no observable enrichment 60 min post-treatment (Fig. 2A, Fig. S2A, Fig. 3A). In contrast to QSK1, the punctate pattern corresponding to CRK2-mVenus peaked at PD only after 60 min of treatment (Fig. 2E). Using QSK1-mCherry and CRK2-mVenus co-expressing lines, we carried out a time-course analysis and showed that, enrichment of QSK1 and CRK2 at PD was sequential (Fig. 3A, Fig. S3A). Following peak enrichment of QSK1-mCherry at PD, CRK2-mVenus began to relocalize to PD. As CRK2-mVenus reached its saturation at PD, we did not observe localization of QSK1-mCherry at PD anymore (Fig. 3A). Overall, we did not observe simultaneous peak enrichment for both QSK1 and CRK2 at PD. This finding suggested that the two RLKs might co-ordinate to precisely fine-tune callose deposition to control the PD aperture.

### CRK2 and QSK1 exist in a pre-formed complex at the bulk PM that dissociates upon osmotic stress

We previously detected QSK1 as an *in vivo* interactor of CRK2^9^ implying that the two RLKs co-localize at the general PM in close proximity by interacting either directly or in a multi-protein complex. However, the sequential recruitment of QSK1 and CRK2 at PD suggested a dissociation of the two upon stress perception. To test this, we performed Förster resonance energy transfer by fluorescence lifetime imaging (FRET-FLIM) of Arabidopsis plants stably co-expressing QSK1-mCherry and CRK2-mVenus proteins (QSK1-mCherry/*crk2* x CRK2-mVenus/*crk2*) with and without mannitol treatment. As a negative control, we used equivalent plants with kinase inactive CRK2 (QSK1-mCherry/*crk2* x CRK2^D450N^-mVenus/*crk2*), since CRK2^D450N^ did not relocalize following stress application^9^. In plants expressing the donor-acceptor pair (CRK2-mVenus + QSK1-mCherry), donor life-time was lower in comparison to the donor-only and negative control plants under control conditions (Fig. 3B, Table S4). This suggested the existence of a preformed complex of CRK2 and QSK1 in normal conditions which would reduce the fluorescence lifetime of the donor mVenus. Interestingly, upon mannitol treatment, the donor lifetime in QSK1-mCherry + CRK2-mVenus plants increased suggesting dissociation of the complex upon mannitol treatment. This finding was in-line with the sequential enrichment of QSK1 and CRK2 at PD (Fig. 3A, Fig. S3A). In contrast, in the QSK1-mCherry + CRK2 ^D450N^-mVenus plants, the donor lifetime was higher under both control and mannitol-treated conditions suggesting that the kinase inactive CRK2 might have weaker affinity for the protein complex formation (Fig. 3B). Further, we performed structural modeling to predict the interaction between CRK2 and QSK1 using Colabfold (version 1.5.2) implementation of AlphaFold2^40,41^. We modeled the full-length proteins, and the combinations of their individual structural segments. The modeled combinations were i) full-length proteins, ii) ExD (Extracellular domains), iii) ExD+TMD (transmembrane domains), iv) TMD, v) TMD+Kinase domain, and vi) Kinase domain for both CRK2 and QSK1. Based on the predicted aligned error (PAE) values (Fig. 3C-D, Fig. S3B) and interface predicted template modelling (iPTM) scores (Table S5), ExD+TMD regions showed high confidence for interaction (Fig. 3C). The modeling of the full-length proteins suggested that the two RLKs engage specific segments for interaction (Fig. 3D) which mainly resulted from the ExD and TMD regions (Fig. S3B). Collectively, these results confirmed that CRK2 and QSK1 are present in close proximity at the general PM and form a complex *in planta,* and this complex dissociates upon osmotic stress perception.

### CRK2 phosphorylates QSK1 on its C-terminus, and salt stress reduces this phosphorylation

The role of QSK1 in PD regulation was reported to be regulated by its phosphorylation^21^ specifically of two serine residues in the C-terminus of the cytosolic region of QSK1 (Ser-621 and Ser-626). However, the responsible kinase was not investigated. Since we previously identified QSK1 as an *in vivo* interactor of CRK2^9^, we asked if CRK2 could phosphorylate QSK1, thereby controlling its plasmodesmal relocalization. We recombinantly expressed the cytosolic parts of CRK2 (6His-GST-CRK2cyto and 6His-GST-CRK2^D450N^cyto)^9^ and QSK1 (6His-MBP-QSK1cyto) in *E. coli*. To identify CRK2-targeted phosphorylation sites in QSK1, we carried out LC-ESI-MS/MS analysis of in-gel trypsin-digested peptides after the kinase reaction of CRK2cyto and QSK1cyto. The identified *in vitro* phosphorylation sites are shown in Table S6 and Fig. 4A. Interestingly, Ser-626 was among the identified phosphosites (Fig. 4A), suggesting that CRK2 was able to directly interact with and phosphorylate the QSK1 peptide containing Ser-626. Due to the importance of Ser-621 and Ser-626 in osmotic stress response^21,32,33,42^, we generated a variant of QSK1 with phospho-negative substitutions at positions 621 and 626 (QSK1^S621A-S626A^) as a phosphorylation target of CRK2. Since 6His-GST-CRK2cyto and 6His-MBP-QSK1cyto have similar molecular weights, we digested 6His-MBP-QSK1cyto and 6His-MBP-QSK1^S621A-S626A^cyto with Factor Xa protease before the kinase reaction to cleave the MBP tag off from the MBP-QSK1cyto fusion protein resulting in a clear resolution of CRK2 and QSK1 cytosolic fragments. The QSK1cyto was phosphorylated by 6His-GST-CRK2cyto but not by the kinase inactive 6His-GST-CRK2^D450N^cyto (Fig. 4B) in *in vitro* kinase assay. Further, we detected an overall comparable phosphorylation signal of the WT QSK1 and QSK1^S621A-S626A^, probably due to multiple other phosphorylation sites present in this part of the protein. (Fig. 4B). Given the role of QSK1 phosphorylation in its relocalization to PD, its *in vitro* phosphorylation by CRK2 suggested a phosphorylation-dependent regulatory function of CRK2 in controlling the subcellular localization of QSK1.

**Figure 4.**
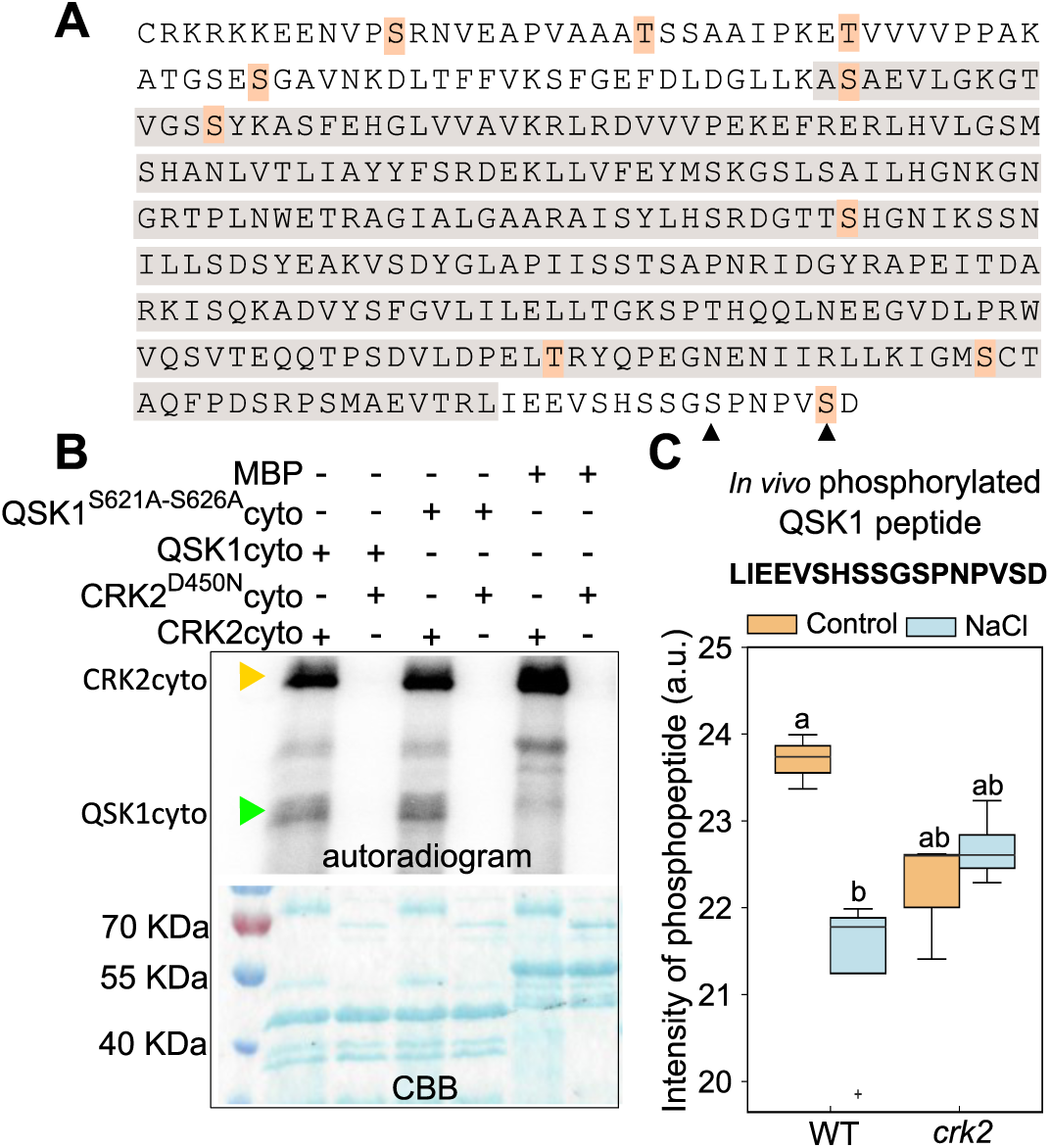
CRK2 phosphorylates QSK1. *In vitro* kinase reaction between the recombinantly expressed cytosolic regions of CRK2 and QSK1. (A) CRK2-targeted phosphosites in the QSK1 cytosolic region as identified by LC-ESI-MS/MS analysis. The highlighted residues in pink are consolidated from 3 experiments-high confidence only. The shown amino acid sequence is the cytosolic region of QSK1 and the grey highlighted part corresponds to the pseudokinase domain. The arrows represent the two serine residues at positions 621 and 626 which were selected for further studies. (B) *In vitro* phosphorylation of QSK1cyto by CRK2cyto. The QSK1cyto-MBP and QSK1^S621A-S626A^ cyto-MBP were digested with Factor Xa protease for 24 h to separate the MBP tag for size resolution before carrying out the kinase reaction. The QSK1 was phosphorylated by CRK2 but not with the kinase inactive CRK2^D450N^. No difference was observed in the phosphorylation of QSK1^S621A-S626A^. The lower blot shows the Coomassie Brilliant Blue (CBB)-stained gel which was destained and developed for the detection of labeled ATP (upper blot). (C) *In vivo* phosphorylation of QSK1. The shown phosphopeptide was detected in the WT and *crk2* seedlings. NaCl treatment and mutation in *CRK2* reduced the intensity of phosphopeptide. The detected phosphopeptide was predicted to be phosphorylated at two serine residues. Boxplot description as Figure 1. n=3-4. Two-way ANOVA, Tukey’s test. See also Table S6 for panel A and Table S7 for panel C.

To determine this *in vivo*, we applied a quantitative phosphoproteomics approach and compared the levels of QSK1 phosphopeptides isolated from WT and *crk2* plants under control and salt stress conditions. We detected the dually phosphorylated peptide LIEEVSHSSGSPNPVSD in both WT and *crk2* under normal and stress conditions. This phosphopeptide contains the Ser-621 and Ser-626 that are previously shown to be phosphorylated in phosphoproteomics studies^42–45^. Interestingly, the abundance of the LIEEVSHSSGSPNPVSD phosphopeptide was reduced following salt treatment in WT but not in *crk2*. Moreover, the abundance of the phosphopeptide was lower in *crk2* compared to WT under normal growth conditions (Fig. 4C, Table S7). Lower abundance in *crk2* may suggest that CRK2 phosphorylates QSK1 in the peptide LIEEVSHSSGSPNPVSD. This finding aligns with the FRET-FLIM assay where we observed that mannitol stress disrupted the binding of CRK2 and QSK1 and placed a role of phosphorylation-dependent coupling of the RLKs in osmotic stress signaling.

### Phosphorylation of QSK1 at serines 621 and 626 inhibits its stress-induced enrichment at PD

QSK1 relocalization in response to osmotic stress is associated with its phosphorylation status at Ser-621 and Ser-626^21^. SIRK1 phosphorylates QSK1 at Ser-621 and Ser-626 during sucrose-induced osmotic conditions^32^. Therefore, we examined if QSK1 relocalization to PD was dependent on SIRK1 by testing QSK1-mCherry/*sirk1* plants. We found that QSK1 exhibited osmotic stress-induced PD enrichment in QSK1-mCherry/*sirk1* plants without impairment. An enrichment index of 1.967 was observed in *sirk1* following osmotic stress compared to 1.092 in control conditions (Fig. S4A-B, Table S3). This finding suggested, that while phosphorylation of Ser-621 and Ser-626 plays an important role in QSK1 recruitment to PD, SIRK1-mediated phosphorylation does not appear to regulate PD recruitment of QSK1.

Since we found that CRK2 also targeted QSK1 phosphorylation in the phosphopeptide LIEEVSHSSGSPNPVSD (Fig. 4) and kinase active CRK2 was required for QSK1 recruitment at PD (Fig. 2C-D), we explored the role of Ser-621 and Ser-626 phosphorylation in context of CRK2. We stably expressed phosphomimetic QSK1 at Ser-621 and Ser-626 by replacing both serines by glutamic acid and aspartic acid in *crk2* (QSK1^(S621D-S626D)^-mCherry/*crk2*; QSK1^(S621E-S626E)^-mCherry/*crk2*) and in WT background as control (QSK1^(S621D-S626D)^-mCherry/WT; QSK1^(S621E-S626E)^-mCherry/WT). We found that enrichment of phosphomimetic QSK1^(S621D-S626D)^-mCherry in the WT background was negligible and statistically insignificant and only showed an increase of enrichment index from 1.045 to 1.119 upon mannitol treatment (Fig. 5A-B). Similarly, negligible relocalization of QSK1^(S621D-S626D)^-mCherry was observed in the *crk2* mutant background (Fig. 5A-B). Consistently, the phosphomimetic variant QSK1^(S621E-S626E)^-mCherry, showed a mild enrichment of 1.336 from 1.017 following mannitol treatment in the WT background and poor enrichment in the *crk2* mutant background (1.011 to 1.117) (Fig. S4C-D). This enrichment of QSK1^(S621E-S626E)^-mCherry in WT background was still weaker and delayed compared to that in the QSK1-mCherry/WT plants. This led us to test the PD recruitment of non-phosphorylatable QSK1, in which the Ser-621 and Ser-626 were replaced by alanines, in WT and *crk2* backgrounds (QSK1^(S621A-S626A)^-mCherry/WT; QSK1^(S621A-S626A)^-mCherry/*crk2*). We observed a higher enrichment of QSK1^(S621A-S626A)^-mCherry at PD in response to mannitol treatment (Fig. 5C-D) in QSK1^(S621A-S626A)^-mCherry/WT plants. To a lesser extent, QSK1^(S621A-S626A)^-mCherry was also present at PD even without mannitol treatment (Fig. 5C-D). The enrichment index for QSK1^(S621A-S626A)^-mCherry/WT in control condition was 1.338 and increased to 2.018 which was even higher than QSK1-mCherry/WT under mannitol treatment (Fig. 2B). We also validated this result in QSK1^(S621A-S626A)^-mCherry/*qsk1* lines and the findings were consistent with a clear and intense enrichment of QSK1^(S621A-S626A)^-mCherry in the *qsk1* mutant background (Fig. S5E-F). These findings confirmed that phosphorylation of QSK1 at Ser-621 and Ser-626 was inhibitory for its recruitment to PD. Further, attenuated or complete inability of QSK1 relocalization to PD in the *crk2* and CRK2^D450N^ backgrounds suggested an essential role for CRK2-mediated phosphorylation of yet unknown factor(s) involved in this process.

**Figure 5.**
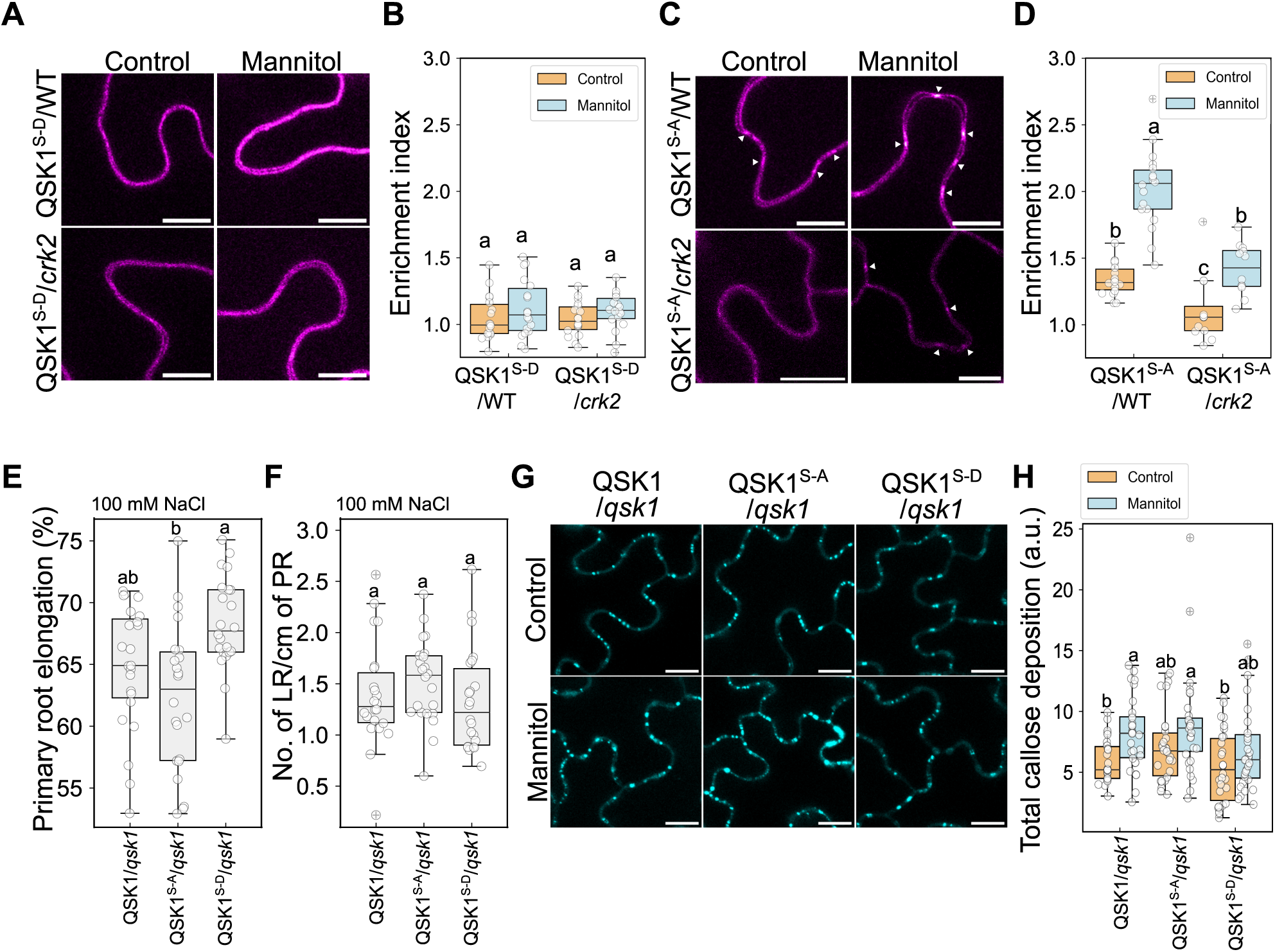
Phosphorylation of QSK1 at Ser-621 and Ser-626 inhibits its recruitment to PD and modulates plant response to osmotic stress. Relocalization of phospho-mutant versions of mCherry-tagged QSK1 in the leaf epidermal cells of Arabidopsis WT or *crk2* mutant backgrounds. (A-B) The phospho-mimetic QSK1^S-D^ (QSK1^(S621D-S626D)^) showed reduced enrichment in both WT and *crk2* mutant backgrounds. (C-D) The phospho-negative QSK1^S-A^ (QSK1^(S621A-S626A)^) showed much higher enrichment in the WT background upon mannitol treatment. A weak enrichment of phospho-negative QSK1 was also observed in control conditions. The phospho-negative QSK1^S-A^ also enriched weakly in the *crk2* mutant background after mannitol treatment. Objective lens 60.0 X oil immersion. Laser transmissivity 7.0%-15%; constant between the comparisons. Experimental conditions and other laser settings for mCherry as Fig. 2. Scale bar 10 µm. Boxplots as described in Figure 1. Two-way ANOVA, Tukey’s test. n ≥ 12. (E-F) Comparison of root response to salt stress. PR elongation is the primary root growth post-treatment. 5-days-old seedlings were transferred to media containing NaCl and were imaged after another 7 days. The PR elongation is calculated respective to the control PR length. n=22-23, two-way ANOVA, Tukey’s test (E). (F) LR density upon NaCl treatment same as for panel E. n=22, Kruskal-Wallis test, Dunn’s multiple comparison test. QSK1, QSK1^S-A^ and QSK1^S-D^ are the plants expressing WT, QSK1^(S621A-S626A)^ and QSK1^(S621D-S626D)^ variant of QSK1, respectively, in the *qsk1* mutant background. (G-H) Visualization and quantification of callose deposition in aniline blue-stained leaf epidermal cells of Arabidopsis. Treatment and imaging as described for Figure 1. Boxplot as described in Figure 1. n = 30, two-way ANOVA, Tukey’s test. See also Fig. S4; Table S3 for panels B, D, Table S8 for panels E-F and Table S9 for panels G-H.

### Phosphorylation of QSK1 at serines 621 and 626 regulates plant stress response and suppresses osmotic stress-induced callose deposition

To test the physiological relevance of QSK1 relocalization to PD and callose deposition during osmotic stress, we carried out root growth assays of plants with WT QSK1 (QSK1-mCherry/*qsk1*), non-phosphorylatable QSK1 (QSK1^(S621A-S626A)^-mCherry/*qsk1*) and phospho-mimetic QSK1 (QSK1^(S621D-S626D)^-mCherry/*qsk1*) with and without salinity stress. While the PR length of all three genotypes was similar under control conditions (Fig. S4I, Table S8), the QSK1^(S621A-S626A)^ plants exhibited maximum retardation in the PR elongation upon salinity stress (Fig. 5E). In contrast, the phospho-mimetic QSK1 plants (QSK1^(S621D-S626D)^) showed even higher PR elongation than the plants with WT QSK1 under salinity stress (Fig. 5E). The LR density, did not statistically vary among the genotypes under both control and salt stress conditions (Fig. S4J, Fig. 5F), nevertheless, a slightly higher LR density was observed in the QSK1^(S621A-S626A)^-mCherry/*qsk1* plants under salt stress and an opposite trend was observed for the QSK1^(S621D-S626D)^-mCherry/*qsk1* plants (Fig. 5F). A positive correlation between QSK1 availability at PD and callose accumulation in the root epidermal layer is already established^21^. Since in our conditions, non-phosphorylatable QSK1 (QSK1^(S621A-S626A)^) was highly enriched at PD in leaf epidermal cells, we tested the effect of QSK1 recruitment at PD on callose accumulation in the leaf epidermis by aniline blue staining. We quantified callose levels after 7 hours of mannitol treatment to QSK1-mCherry/*qsk1*, QSK1^(S621A-S626A)^-mCherry/*qsk1* and QSK1^(S621D-S626D)^-mCherry/*qsk1* plants (Fig. 5G-H, Fig. S4G-H, Table S9). Plants expressing WT or non-phosphorylatable QSK1 (QSK1^(S621A-S626A)^) accumulated higher callose upon mannitol treatment compared to their respective control conditions. However, plants expressing phosphomimetic QSK1 (QSK1^(S621D-S626D)^) responded weakly to mannitol treatment (Fig. 5G-H, Fig. S4G-H). In control conditions, slightly higher callose levels were detected in the QSK1^(S621A-S626A)^ plants in terms of both intensity and number of deposits compared to the plants with WT QSK1 (Fig. 5G-H, Fig. S4G-H) which is consistent with a higher basal presence of QSK1^(S621A-S626A)^ at PD even without stress treatment (Fig. 5C-D), however, the differences in the overall basal callose accumulation were statistically insignificant among the three genotypes which is consistent with our finding that QSK1 might not play a regulatory role in basal callose deposition (Fig. 1E-F). Collectively, these results established the physiological relevance of the QSK1 relocalization to PD during osmotic stress and highlighted protein phosphorylation as a regulatory mechanism in growth adjustments.

## Discussion

Symplastic communication through PD is crucial for coordinating plant growth and stress responses^14,15,46–48^. Dynamic regulation of symplastic connectivity is achieved through PD-associated proteins^49^, lipid composition^50^ and callose deposition^18^. Current knowledge on the changes of callose accumulation and symplastic communication is largely focused on immune response^19,20,51–53^. Studies show altered symplastic communication during abiotic stress conditions such as metal toxicity^54,55^, low temperature^56^ and mechanical wounding^57^. Despite the widespread occurrence of osmotic stress in plants, understanding of how callose accumulation and PD connectivity respond to it remains limited^9–11,21^.

RLKs are involved in sensing and integrating extracellular signals and they typically function in complexes rather than acting alone. For example, FLS2-BAK1, EFR-BAK1 and CERK1-LYK5 (Chitin elicitor receptor kinase 1-Lysin motif-containing receptor-like kinase 5)^58^ in immune signaling^59,60^, BRI1-BAK1 in BR signaling^61^, SIRK1-QSK1 in regulation of aquaporins^32,33^. Importantly, the combination of different RLKs can determine the functional outcome. The RLKs CRK2 and QSK1 mediate plant growth and osmotic stress response through plasmodesmal regulation^9,21^, however, how they coordinate to control callose levels during osmotic stress is so far not understood. In this study, we identify a regulatory mechanism that precisely fine-tunes callose deposition during osmotic stress response, a key part of plant growth adaptation to changing environmental conditions. We report that under normal growth conditions, CRK2 inhibits callose deposition by engaging QSK1 at the general PM where the CRK2-QSK1 complex potentially has another role specific to the general PM.

The growth phenotypes of *crk2*, *qsk1* and the *crk2 qsk1* mutants suggested that CRK2 and QSK1 are important for normal plant growth and stress response. The *crk2* mutant exhibits strongest phenotypic alterations compared to WT Arabidopsis which may be due to pleotropic functions of CRK2^9,37,62,63^. The smaller rosette and salt hypersensitivity during germination in the *crk2* mutant was rescued in the *crk2 qsk1* mutant suggesting antagonistic roles of CRK2 and QSK1 in the biological processes that result in observed phenotypes. Further, mutants defective in callose regulation have defects in LR patterning and PR length due to altered symplastic connection^9,12,17^. In our analyses, the root phenotype of the *crk2 qsk1* mutant was milder compared to the *crk2* and did not rescue to the WT level implying that the absence of both functional CRK2 and QSK1 results in defects in processes regulating PR growth and LR emergence.

Further, CRK2 and QSK1 also showed antagonistic effects on callose deposition especially under osmotic stress conditions. A higher callose level in *crk2* suggested an inhibitory role in callose deposition which could result from multiple yet unknown actions of CRK2. For example, CRK2-targeted phosphorylation of CALLOSE SYNTHASES (CALS)^9^ could be one of the regulatory mechanisms determining callose levels. Further, CRK2 also regulates ROS production through RBOHD^62^ which could add another layer of CRK2-mediated regulation of callose production. In the *qsk1* mutant, a reduced callose deposition upon mannitol treatment suggested that the inhibitory action of CRK2 is ensued following osmotic stress despite the absence of QSK1 which gets further support from the result that QSK1 was not found to be necessary to recruit CRK2 at the PD. In the *crk2 qsk1* mutant, the WT-like response indicated alleviation of the inhibitory effect of CRK2 and QSK1 on each other and possibly the presence of compensatory or feedback mechanisms that maintain callose homeostasis.

Protein phosphorylation is a common regulatory mechanism across living organisms for dynamically reshaping the binding repertoire of a given protein and thereby providing functional specificity^64,65^. The changes in the protein-protein interaction networks by phosphorylation can also lead to changes in the localization and protein abundance^30,66^. A regulatory role of QSK1 phosphorylation at Ser-621 and S-626 has been previously describd^42–44^. For example, QSK1 phosphorylation at Ser-621 and Ser-626 by SIRK1, an LRR RLK, stabilizes the SIRK1-QSK1 protein complex involved in the regulation of aquaporins in roots^32^. QSK1 phosphorylation at these two residues also enhances its interaction with the nitrate transceptor NRT1.1 which eventually transduces low nitrate signals to PM H^+^-ATPase AHA2^36^.

Through *in vitro* kinase assays, we identified that CRK2 phosphorylates multiple residues in the cytosolic region of QSK1 including Ser-626. Further, *in vivo* phosphorylation status of QSK1 within LIEEVSHSSGSPNPVSD peptide was altered in the *crk2* mutant. QSK1 was shown to interact with CRK2 *in planta*^9^, and FRET-FLIM analyses together with structural modeling supported that CRK2 and QSK1 exist in a preformed complex at the general PM (Fig. 6 panel A). Interestingly, the CRK2 and QSK1 affinity was abolished when CRK2 was replaced by CRK2^D450N^. Together, these results and existing literature suggested that phosphorylation of QSK1 at Ser-621 and Ser-626 could be a common regulatory mechanism to stabilize QSK1 protein complexes, including the CRK2-QSK1 complex. *In vivo* phosphoproteomics analyses suggested reduced phosphorylation of the LIEEVSHSSGSPNPVSD peptide at two serine residues following salt treatment in WT. Consistently, mannitol treatment reduced the binding affinity of the CRK2-QSK1 complex in the FRET-FLIM assays. These findings suggest, that an unknown protein phosphatase might act on QSK1 to allow dissociation of the CRK2-QSK1 complex (Fig. 6 panel B) leading to the PD localization of QSK1 prior to the relocalization of CRK2. Consistently, our microscopy-based relocalization studies showed that phosphorylation of QSK1 at Ser-621 and Ser-626 is inhibitory for the recruitment of QSK1 to PD and consequently callose accumulation following mannitol treatment. Non-phosphorylatable QSK1^S621A-S626A^, efficiently relocalized to PD while QSK1 phosphomimetic substitutions at Ser-621 and Ser-626 abated its recruitment to PD. Collectively, these results demonstrated a phosphorylation-dependent subcellular dichotomy for QSK1 conditionally associating with and dissociating from CRK2 and probably other proteins. This mechanism could direct QSK1 towards distinct molecular pathways to promote growth adaptability under osmotic stress.

**Figure 6.**
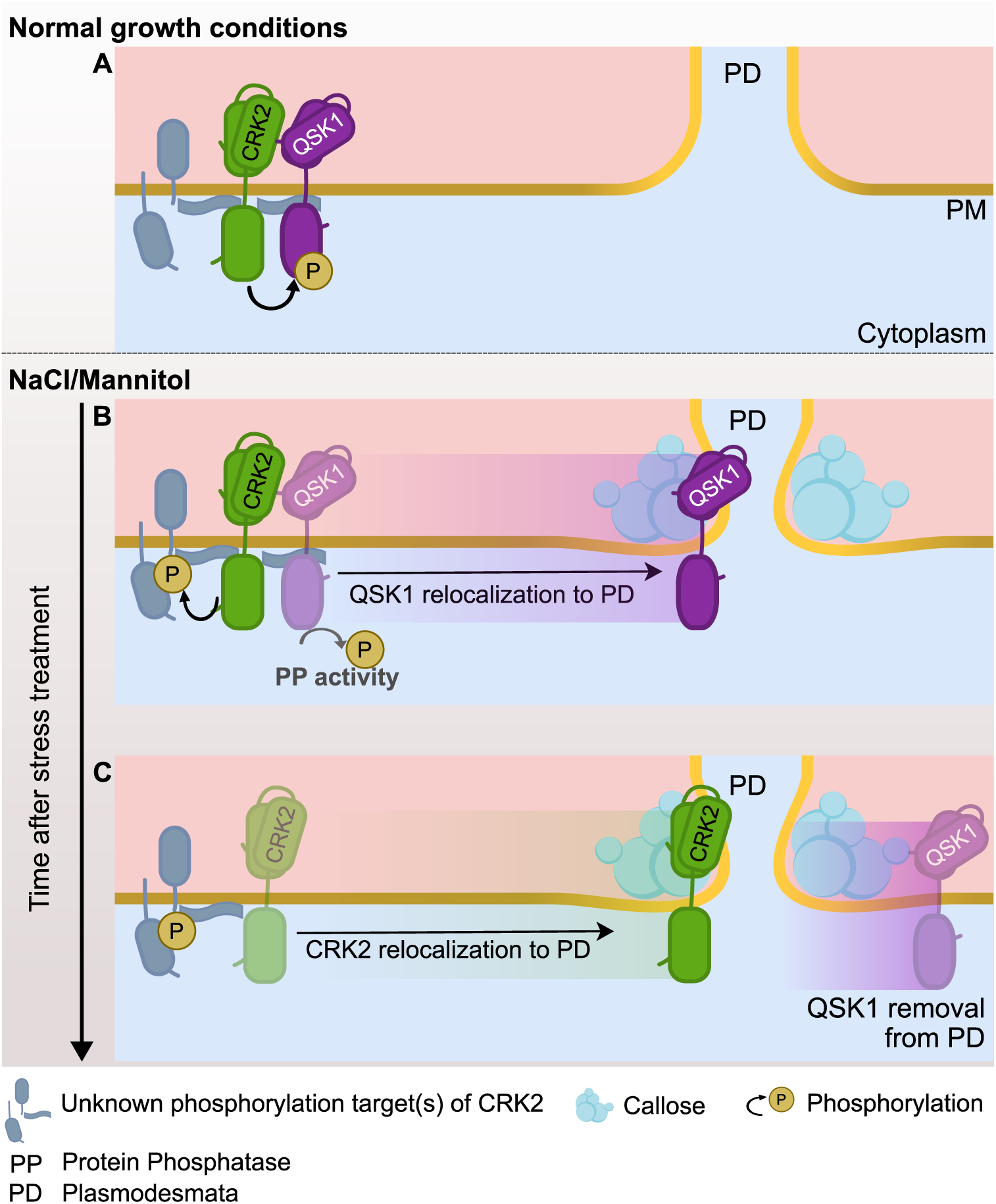
Proposed model- CRK2 modulates plasmodesmal function of QSK1. Under normal growth conditions, CRK2 phosphorylates QSK1 and the two RLKs form a complex at the bulk plasma membrane (PM) inhibiting unnecessary recruitment of QSK1 to PD (A). Exposure to salt or mannitol promotes an unknown protein phosphatase activity at QSK1 releasing it from the CRK2-QSK1 complex and consequently QSK1 relocalization to PD. CRK2 also phosphorylates another unknown protein(s), a necessary event for QSK1 enrichment at PD. QSK1 positively regulates callose deposition at PD (B). Subsequently, QSK1 is removed from PD and returns to basal level. CRK2 enrichment at PD follows (C).

Although phosphorylation of QSK1 was inhibitory for its relocalization to PD (Fig. 5A-D), enzymatically active CRK2 was still essential for QSK1 recruitment at PD (Fig. 2A-D). This suggested the possibility that CRK2 might phosphorylate another protein(s) in addition to QSK1, whose phosphorylation by CRK2 would be necessary to facilitate the relocalization of QSK1 to PD (Fig. 6 panel B). The PM could be biochemically divided into detergent resistant membrane (DRM) and detergent soluble membrane (DSM) domains, and PD-associated membranes are considered to be DRMs. Proteins identified in PD proteomes and DRM fractions show a large overlap^21^ and QSK1 has been found in several DRM and PD proteomes (compiled in^21^). The actin cytoskeleton is necessary for the distribution of proteins between DRM and DSM and disruption of the actin cytoskeleton depletes QSK1 from DRMs^67^. We previously identified actin isoforms, as well as other DRM-localized proteins as interactors of CRK2 in pull-down assays followed by MS analyses^9^. Therefore, we speculate on regulation of the actin cytoskeleton or other DRM-localized proteins by CRK2 to enable QSK1 relocalization after its dephosphorylation, however, more studies are required to support this.

Adjusting root system architecture is one of the main strategies plants employ during extended periods of salinity stress for efficient water uptake^68^. QSK1 phosphorylation at Ser-621 and Ser-626 affects callose deposition in the epidermal layer of root ^21^ as well as leaves, resulting in altered symplastic connectivity and root phenotype. Adjusting callose levels through QSK1 can therefore be a contributory mechanism plants employ for osmotic stress resilience. In our physiological analyses, we found that the plants accumulating higher callose levels upon osmotic stress, exhibit a root phenotype adjusted for osmotic stress.

Based on these findings, we propose a working model in which symplastic connectivity is dynamically regulated through temporally coordinated signaling between CRK2 and QSK1 (Fig. 6). We propose that CRK2 works as a moderator of QSK1-mediated osmotic stress responses. CRK2 phosphorylates QSK1 and sequesters it at the general PM. This complex might have a function specific to their loacalization at the general PM. At the same time, existence of such a complex inhibits unnecessary QSK1 function at PD in the absence of osmotic stress. Dephosphorylation of QSK1 at the C-terminus upon stress stimuli dissociates it from CRK2 allowing it to relocalize to PD to positively regulate callose deposition. At a later stage, CRK2 moves to PD itself where it likely finds different phosphorylation targets than at the PM and eventually negatively regulates callose deposition to switch it off and prevent unnecessary clogging of PD. In conclusion, our work shows that CRK2 controls the dynamic and stimulus-dependent relocalization of QSK1, and likely also other proteins, to PD. PD and other membrane nanodomains contain constitutive protein components^23^, however, the dynamics of QSK1 and CRK2 demonstrate the necessity and potential for the precise spatiotemporal regulation of the cellular response machinery. In the future, it will be highly interesting to investigate the spatiotemporal dynamics of other proteins showing conditional clustering in nanodomains vs. the general plasma membrane.

## Material and methods

### Plant materials and growth conditions

The *Arabidopsis thaliana* ecotype Col-0 was used as wild-type control. The seeds of mutants *qsk1* (SALK_019840; AT3G02880), *crk2* (SALK_012659C; At1g70520), and *sirk1* (SALK_125543C; At5g10020) were obtained from NASC (University of Nottingham). The *crk2 qsk1* double mutant was generated by crossing *qsk1* and *crk2*. For growing plants under sterile conditions, seeds were surface-sterilized and stratified in dark for 3 days at 4°C and plated on 0.5X Murashige and Skoog media (Sigma-Aldrich) supplemented with 0.8% (w/v) agar, 1% (w/v) Suc, and 0.1% (w/v) MES, pH 5.8. This media composition was used as standard growth media for all the *in vitro* experiments unless specified. Plants were grown in climate-controlled growth chambers with a 16-h light, 8-h dark photoperiod. For experiments with adult stages, seed propagation, and growth of selected transgenics, the plants were grown in substrate mix (2:1 peat:vermiculite) in a Weiss Gallenkamp growth chamber or greenhouse with long-day conditions at 21°C.

### Physiological assays

For comparative phenotypic analysis of WT, *crk2*, *qsk1*, and *crk2 qsk1* plants, 10 plants for each genotype were planted in individual pots with substrate mix (2:1 peat:vermiculite). Plants were allowed to grow in a Weiss Gallenkamp growth chamber with long-day conditions at 21°C with equal irrigation and randomization of pot positioning. Images were captured at various growth stages. Related data and statistical analysis are given in Table S1.

For root elongation assays, 5-day-old seedlings grown in vertically positioned square petri plates in Photon Systems Instruments (PSI) growth chamber were transferred to standard 0.5X MS medium supplemented with 100 mM NaCl and the primary root tip was marked. For the control group, the seedlings were transferred to medium without NaCl. The images were captured after another 7 days and the root elongation was measured with the program Fiji (https://imagej.net/software/fiji/). The data obtained were the average of at least 13-15 seedlings and all the experiments were repeated three times unless specified. Related data and statistical analysis are given in Table S1.

For the seed germination assay, surface-sterilized seeds were plated evenly in petri dishes with standard 0.5X MS medium supplemented with 150 mM NaCl. For the control group, the seeds were sown on the medium without NaCl. The seedlings were allowed to grow in a PSI growth chamber and the images of the plates were captured after 12 days. For each genotype and treatment, 3 plates were taken as biological replicates and in each plate 30-35 seedlings were taken. The percentage of seed germination was first estimated in control plates to normalize the effect of non-viable seeds. The total number of seeds and the viable germinated seeds with clearly visible root and green cotyledon/leaves were counted. Related data and statistical analysis are given in Table S1.

### Osmotic stress treatment and callose estimation

For estimation of osmotic stress-induced callose deposition, seeds were sown hydroponically for two weeks on a mesh in liquid 0.5X MS with 1% sucrose in standard growth conditions. For inducing callose deposition under osmotic stress, the roots were subjected to 50 mM mannitol in liquid medium for 7 hours. After treatment, the plants were pulled out of the mesh and infiltrated with 0.2%-0.5% aniline blue (constant for a single experiment) and the first pair of true leaves were imaged in a laser confocal microscope. For each genotype, 5 plants were imaged and 5-6 areas were scanned from first pair of true leaves of each plant. The control plants underwent the same process without mannitol treatment. The mean intensity and number of deposits were measured using Fiji. To quantify overall callose levels that include both intensity and number of deposits, a composite metric was derived by multiplying mean intensity of deposits and number of deposits and dividing it with a constant scaling factor to yield a number suitable for comparative analysis. This is termed as total callose deposition. Related data and statistical analysis are given in Table S2 and S9.

### Plasmid vector construction and transgenic lines

Plasmid constructs for P*_QSK1_*:*QSK1*-mCherry, P*_QSK1_*:*QSK1*^(S621A-S626A)^-mCherry, P*_QSK1_*:*QSK1*^(S621D-S626D)^-mCherry and P*_QSK1_*:*QSK1*^(S621E-S626E)^-mCherry were generated by the MultiSite Gateway technology (Invitrogen; Thermo Fisher Scientific). The donor clones for promoter regions of *QSK1* and coding sequence of *QSK1* were cloned in the vectors pDONR™P4-P1R and pDONR™221, respectively. The donor clone for mCherry in the vector pDONR™P2R-P3 was already available in the laboratory. The donor clones were combined in an LR reaction to obtain the final constructs in the binary vector pH7m34GW (VIB-UGENT). The phospho-mutant versions of QSK1 (S621A-S626A, S621D-S626D and S621E-S626E) were generated by introducing point mutation by DpnI-mediated site-directed mutagenesis^70^. The primers are listed in Table S10. P*_CRK2_*:*CRK2*- mVenus and P*_CRK2_*:*CRK2^D^*^450^*^N^*-mVenus constructs were available in the binary vector pBm43GW (Invitrogen; Thermo Fisher Scientific) (Hunter et al 2019). The final clones in the binary vectors were introduced in *Agrobacterium tumefaciens* GV3101_pSoup and Arabidopsis plants of desired genotypes were transformed by floral dip method^71^. The CRK2/CRK2 *^D^*^450^*^N^*-mVenus and QSK1-mCherry co-expressing lines were generated by crossing P*_CRK2_*:*CRK2*-mVenus/*crk2* and P*_QSK1_*:*QSK1*-mCherry/*crk2*, and, P*_CRK2_*:*CRK2^D^*^450^*^N^*-mVenus/*crk2* and P*_QSK1_*:*QSK1*-mCherry/*crk2* plants. Transformants were selected on 0.5X MS medium with appropriate antibiotic or herbicide [(hygromycin 20 ug/ml (CARL ROTH) or Basta 10 ug/ml (DL-phosphinothricin; Duchefa Biochemie)].

### Osmotic stress treatment and confocal imaging

For visualizing localization of the fluorescently-tagged proteins in response to osmotic stress, the mature leaves of 4-week-old Arabidopsis were detached from the plant and submerged in 0.4 M mannitol or 150 mM NaCl in 0.5X MS solution. For detecting co-localization with aniline blue, the samples were treated with 0.2% aniline blue simultaneously with mock or 0.4 M mannitol and 150 mM NaCl treatments. After the incubation period, the same area of all the leaf samples was targeted for imaging and mounted in the same solution and the abaxial side was imaged. A mock-treated leaf of the same plant was used as a control. All the live-imaging experiments were conducted on an Olympus FLUOVIEW FV3000 confocal laser scanning microscope with 60.0 X or 40.0 X oil-immersion objectives and was operated by Olympus FV31S-SW software. The excitation wavelengths for mCherry, mVenus and DAPI (for aniline blue) were 561, 488 and 405, respectively. The detection wavelengths were 570-620 nm for mCherry, 500-540 nm for mVenus and 450-550 nm for DAPI. The laser transmittivity was kept constant for a single experiment and for the comparisons of fluorescence intensities unless otherwise specified. Co-localizations were detected in sequential scanning mode to prevent cross bleeding of emission spectra.

### Quantification of protein enrichment

To quantify the extent of protein enrichment, we used the method described in ^72^ with modification. We assumed that the intense dots of the fluorescently-tagged protein correspond to PD based on the fact that these dots co-localized with the aniline blue-stained callose dots in our analysis (Fig. S2B). Under salt and mannitol treatments, the punctate pattern of CRK2 and QSK1 correspond to their PD localization and previous studies have confirmed this with co-localization with PD-located protein markers (PDLPs) in addition to aniline blue^9,21^. Using Fiji, we selected a circular region of interest (ROI) on such intensity spots resembling PD. The ROI adjacent to the enrichment spots was considered as bulk PM and the ratio of the two was referred to as ‘enrichment index’. In each image, 5-6 pairs of such ROIs were measured for the intensities and enrichment indexes were determined. 3 images for each treatment/genotype were analyzed. The relocalization experiments were conducted on 3 independent transgenic lines for each genotype. The data related to enrichment index and statistical analysis are given in Table S3.

### FRET-FLIM

The experiments on F rster resonance energy transfer detected by fluorescence lifetime (FRET-FLIM) were performed on the confocal microscope FV3000 (Olympus) equipped with the 40.0 X UApo N water immersion objective lens NA 1.15 (Olympus), MultiHarp 150 Time-Correlated Single Photon Counting device, pulsed 488 nm laser, and two Hybrid Photomultiplier detectors (all Picoquant). All data were acquired at 40 MHz pulse repetition rate. Emitted fluorescence was filtered using the FF02-525/40-25 emission filter (Semrock). Data analysis was done using Picoquant SymphoTime64 software package. Membrane regions were manually selected in the images, and lifetime (τ) values were calculated from decay curves using exponential tailfit with the fixed time window for analysis. The acceptable goodness of fit χ2 values were between 0.9 and 1.2. The stage of the plants and treatment used in FRET-FLIM analysis are as described in the method section ‘osmotic stress treatment and live confocal imaging’. Related data and statistical analysis are given in Table S4.

### Structural modeling

The interactions of full-length and truncated CRK2 and QSK1 were predicted using Colabfold (version 1.5.2) implementation of AlphaFold2^40^ multimer version 1 with a maximum of 48 recycles or 0.5 tolerance. The amino acid boundaries selected for both the proteins are given in Table S5.

### Identification of phosphopeptides from *in vitro* kinase assay

The cytosolic region of QSK1 coding sequence QSK1cyto was amplified and cloned into pMal-c2X (Addgene, #75286) using *BamH*I and *Sal*I restriction sites. The primers are given in Table S10. The phospho-negative QSK1cyto(S621A-S626A) was generated by *Dpn*I-mediated site-directed mutagenesis and cloned in the same way as for the WT construct. The construct for the cytosolic region of CRK2 was available (CRK2cyto-pOPINM;^62^). GST-tagged CRK2cyto and MBP-tagged QSK1cyto were expressed in the *E. coli* strain SHuffle T7 (New England Biolabs, C3026J) and BL21-CodonPlus-RIL (Agilent, 230240), respectively, and eventually purified with glutathione sepharose 4B (GE Healthcare, 17-0756-01) and amylose resin (New England Biolabs E8021S) beads, respectively. 10mM maltose was used for the elution of MBP-tagged proteins, while 20mM reduced glutathione for the elution of GST-tagged proteins. In vitro kinase assay was performed as described in Kimura et al. 2020, with minor modifications. Briefly, expressed and purified recombinant proteins were incubated in the ratio 1: 3-5 (kinase: substrate) for 30 min at room temperature in reaction buffer containing γ-32P labelled ATP (0.2µl/20µl of labelled ATP with declared activity 9.25MBq), 50mM HEPES (pH 7.4), 1mM DTT, 10mM MgCl2, 60µM unlabeled ATP. Any specific procedure applied to a specific Figure panel is given in its Figure legend. Enzymatic reactions were terminated by boiling the samples in protein loading dye (98°C, 5 minutes) and proteins were separated by SDS-PAGE. Gels were stained with Coomassie Brilliant Blue and dried. Dried gels were exposed to photosensitive layer and documented with imaging system Typhoon 9410 (Molecular Dynamics). For the purpose of the identification of phosphorylation sites, labelled ATP was replaced by the respective volume of deionized water. Separation and staining of the gels were followed by in-gel trypsin digestion (Thermo Fisher Scientific, 90057). Digested peptide samples were dissolved in 10 µl of 0.1% formic acid and 5 µl was injected for LC-ESI-MS/MS analysis. The LC-ESI-MS/MS analysis was performed on a nanoflow HPLC system (Easy-nLC1000, Thermo Fisher Scientific) coupled to the Q Exactive HF mass spectrometer (Thermo Scientific, Bremen, Germany) equipped with a nano-electrospray ionization source. Peptides were first loaded on a trapping column and subsequently separated inline on a 15 cm C18 column (75 μm x 15 cm, ReproSil-Pur 3 µm 120 Å C18-AQ, Dr. Maisch HPLC GmbH, Ammerbuch-Entringen, Germany). The mobile phase consisted of water with 0.1% formic acid (solvent A) and acetonitrile/water (80:20 (v/v)) with 0.1% formic acid (solvent B). Peptides were eluted using a 30 min gradient: 5% to 21% solvent B in 11 min, 21% to 36% solvent B in 9 min, and 36% to 100% solvent B in 5 min, followed by a 5 min wash with 100% solvent B. MS data was acquired automatically by using Thermo Xcalibur 4.1 software (Thermo Scientific). The data-dependent acquisition method consisted of an Orbitrap MS survey scan of mass range 350–1750 m/z with a resolution of 120000 and AGC target of 3000000. This was followed by HCD fragmentation for 10 most intense peptide ions with resolution of 15000 and AGC target of 5000. Raw files were searched for protein identifications using Proteome Discoverer 2.5 software (Thermo Fisher Scientific) connected to an in-house Mascot 2.8.2 server (Matrix Science). Data was searched against a SwissProt database (version 2022_03) with taxonomy restricted to *Arabidopsis thaliana*.

### Phosphoproteomics

#### Plant treatment

Arabidopsis seeds were sterilized by incubating with 1.5% NaClO 0.02% Triton X-100 solution for 5 min and vernalized at 4 °C for 2 days. Sterilized seeds were germinated and grown in liquid culture on 12 well plates (30 seeds/well) in 1/2 MS medium with 1% sucrose and 0.5g MES/l at 23 °C under continuous light (100 µmol m^-2^ s^-1^) in a Percival growth chamber. Plates with 7-day old seedlings were transferred from the growth chamber to a workbench and kept o/n for acclimatization before treatments. Seedlings were treated with either 150 mM NaCl or sterile water for 30 minutes after which seedlings were immediately collected and flash-frozen in liquid nitrogen and stored at -80 °C.

#### Sample preparation and phosphopeptide enrichment

Frozen seedlings were disrupted using a Retsch mill with 2 steel beads at 50 Hz for 5 min. To the finely ground material was added 500 µL urea extraction buffer (8M in 100 mM Tris-HCl pH 8.5, 20 µL/mL phosphatase inhibitor cocktail 2 (Sigma, P5726), 20 µL/mL phosphatase inhibitor cocktail 3 (Sigma, P0044), 5 mM DDT), samples were vortexed thoroughly to ensure wetting of all powder while thawing. Samples were incubated at RT for 30 min with shaking. Next samples were centrifuged for 5 min at 10k x g, supernatants were transferred to fresh tubes and centrifuged at full speed for 10 mins. This procedure was repeated until no visible pellet was retained. Protein concentration of the extract was determined using Pierce 660 mn protein assay. Samples were alkylated using chloroacetamide (CAA) (550 mM stock, 14 mM final concentration) for 30 min at RT in the dark, after which samples were subjected to an in-solution digestion. In brief, samples (400 µL) were diluted with 400 µL 100mM Tris, pH 8.5, 1mM CaCl_2_, then 5 µg Lys-C (Merck) was added. Samples were incubated for 4h at room temperature, after which 2.4 mL 100mM Tris, pH 8.5, 1 mM CaCl_2_ and 5 µg trypsin were added. Samples were incubated o/n at 37 °C. Samples were acidified with 20 µL TFA and desalted using C18 SepPaks (1cc cartridge, 100 mg (WAT023590)). In brief, SepPaks were conditioned using methanol (1 mL), buffer B (80% acetonitrile, 0.1% TFA) (1 mL) and buffer A (0.1% TFA) (2 mL). Samples were loaded by gravity flow, washed with buffer A (1 x 1 mL, 1x 2 mL) and eluted with buffer B (2 x 400 µL). 40 µL of eluates were used for peptide measurement and concentrated as input control. For phosphopeptide enrichment by metal-oxide chromatography (MOC) (adapted from: Nakagami 2014) the samples were evaporated to a sample volume of 50 µL and diluted with sample buffer (2 mL AcN, 820 µL lactic acid (LA), 2.5 µL TFA / 80% ACN, 0.1% TFA, 300mg/ml LA, final concentrations) (282 µL). MOC tips were prepared by loading 100 µL of a well-mixed slurry of 15mg metal oxide beads (Titansphere TiO_2_ beads 10 µm (GL Science Inc, Japan, Cat. No. 5020-75010) in 500 µL MeOH onto 4 C2 micro columns/condition (3 mg/column) and centrifugation for 5 min at 1500g. Tips were washed with centrifugation at 1500g for 5 min using 80 µL of solution B (80% acetonitrile, 0.1% TFA) and 80 µL of solution C (300 mg/mL LA in solution B). To simplify the processing, samples tips were fitted onto a 96/500 µL deep well plate (Protein LoBind, (Eppendorf Cat. No. 0030504100). After washing MOC tips were transferred to a fresh plate, samples were loaded onto the equilibrated tips and centrifuged for 10 min at 1000g. The flow through was reloaded onto the tips and centrifugation was repeated. Tips were washed with centrifugation at 1500g for 5 min using 80 µL of solution C and 3x 80 µL of solution B. For the elution of the enriched phosphopeptides the tips were transferred to a fresh 96/500 µL deep well plate containing 100 µL/well of acidification buffer (20% phosphoric acid). Peptides were eluted first with 50 µL elution buffer 1 (5% NH_4_OH) and centrifugation for 5 min at 800g, then with 50 µL of elution buffer 2 (10% piperidine) and centrifugation for 5 min at 800g. Next, the samples were desalted using StageTips with C18 Empore disk membranes (3 M) ^74^, dried in a vacuum evaporator, and dissolved in 10 µL 2% ACN, 0.1% TFA (A* buffer) for MS analysis. ***Data acquisition-*** Samples were analysed using an Ultimate 3000 RSLC nano (Thermo Fisher) coupled to an Orbitrap Exploris 480 mass spectrometer equipped with a FAIMS Pro interface for Field asymmetric ion mobility separation (Thermo Fisher). Peptides were pre-concentrated on an Acclaim PepMap 100 pre-column (75 µM x 2 cm, C18, 3 µM, 100 Å, Thermo Fisher) using the loading pump and buffer A** (water, 0.1% TFA) with a flow of 7 µl/min for 5 min. Peptides were separated on 16 cm frit-less silica emitters (New Objective, 75 µm inner diameter), packed in-house with reversed-phase ReproSil-Pur C18 AQ 1.9 µm resin (Dr. Maisch). Peptides were loaded on the column and eluted for 130 min using a segmented linear gradient of 5% to 95% solvent B (0 min: 5%B; 0-5 min -> 5%B; 5-65 min -> 20%B; 65-90 min ->35%B; 90-100 min -> 55%; 100-105 min ->95%, 105-115 min ->95%, 115-115.1 min -> 5%, 115.1-130 min ->5%) (solvent A 0% ACN, 0.1% FA; solvent B 80% ACN, 0.1%FA) at a flow rate of 300 nL/min. Mass spectra were acquired in data-dependent acquisition mode with a TOP_S method using a cycle time of 2 seconds. For field asymmetric ion mobility separation (FAIMS) two compensation voltages (-45 and -65) were applied, the cycle time for the CV-45 experiment was set to 1.2 seconds and for the CV-65 experiment to 0.8 sec. MS spectra were acquired in the Orbitrap analyzer with a mass range of 320–1200 m/z at a resolution of 60,000 FWHM and a normalized AGC target of 300%. Precursors were filtered using the MIPS option (MIPS mode = peptide), the intensity threshold was set to 5000, Precursors were selected with an isolation window of 1.6 m/z. HCD fragmentation was performed at a normalized collision energy of 30%. MS/MS spectra were acquired with a target value of 75% ions at a resolution of 15,000 FWHM, at an injection time of 120 ms and a fixed first mass of m/z 120. Peptides with a charge of +1, greater than 6, or with unassigned charge state were excluded from fragmentation for MS^2^.

#### Data analysis phosphoproteome analysis

Raw data were processed using MaxQuant software (version 1.6.3.4, http://www.maxquant.org/)^75^ with label-free quantification (LFQ) and iBAQ enabled^76^. MS/MS spectra were searched by the Andromeda search engine against a combined database containing the sequences from *A. thaliana* (TAIR10_pep_20101214 database), and sequences of 248 common contaminant proteins and decoy sequences. Trypsin specificity was required and a maximum of two missed cleavages allowed. Minimal peptide length was set to seven amino acids. Carbamidomethylation of cysteine residues was set as fixed, phosphorylation of serine, threonine and tyrosine, oxidation of methionine and protein N-terminal acetylation as variable modifications. The match between runs option was enabled. Peptide-spectrum-matches and proteins were retained if they were below a false discovery rate of 1% in both cases. Statistical analysis of the intensity values obtained for the phospho-modified peptides (“modificationSpecificPeptides.txt” output file) was carried out using Perseus (version 1.5.8.5, http://www.maxquant.org/). Intensities were filtered for reverse and contaminant hits and the data was filtered to retain only phospho-modified peptides. Next. intensity values were log2 transformed. After grouping samples by condition only those sites were retained for the subsequent analysis that had three valid values in one of the conditions. Two-sample t-tests were performed using a permutation-based FDR of 0.05. Alternatively, the data was filtered for 4 valid values in one of the conditions, the valid value-filtered data was median-normalized and missing values were imputed from a normal distribution, using the default settings in Perseus (1.8 downshift, separately for each column). Volcano plots were generated in Perseus using an FDR of 5% an S0=1. The Perseus output was exported and further processed using Excel.

### Quantification and statistical analysis

Fiji (ImageJ; (https://imagej.net/software/fiji/) was used for image analyses. The information on statistical analyses is given in the corresponding figure legends and supplementary tables. Instant Clue^77^ and GraphPad Prism v10 were used for graph preparation and statistical analyses. In general, data normality was tested by Shapiro-Wilk test. Kruskal-Wallis test followed by Dunn’s multiple comparison test was used for those datasets that did not pass the normality test. For normal data, one-way ANOVA with Tukey’s post-hoc test was used. For the datasets with two variables (genotype and treatment), two-way ANOVA was applied with Tukey’s post-hoc multiple comparison test. Any other specific tests are mentioned in figure legends and corresponding supplementary tables.

## Resource availability Contact

Requests for further information and resources in this study should be directed to and will be fulfilled by the lead contact, Michael Wrzaczek.

## Materials availability

Plasmids and transgenic plant seeds generated in this study are available from the lead contact upon request.

## Data and code availability

- This paper does not report original code.
- The raw MS data from this study (phosphoproteomics profiling) was deposited at the ProteomeXchange Consortium via the PRIDE^69^ partner repository with the dataset identifier PXD070133.

## Acknowledgments

This work was supported by the Czech Science Foundation grants nr. 23-04866S (MW) and 22-35680M (RP). FC was funded by EMBO (award number: ALTF 1115-2021; 2022-2024) and the Czech Ministry of Youth and Education MSCA Fellowships Interfellows (Project number: CZ.02.01.01/00/22_010/0003414; 2024-2025). We acknowledge Prof. Jiří Friml and Dmitrii Vladimirtsev (Institute of Science and Technology, Austria) for assistance with FRET-FLIM. We thank Laboratory of Microscopy and Histology, Biology Centre CAS, supported by project LM2023050 funded by the Ministry of Education, Youth and Sports of the Czech Republic, with the instrumental equipment co-financed by the European Union. We also acknowledge Growth Facility (Jan Kadlec), BC Core Facilities, Institute for Plant Molecular Biology, Biology Centre CAS. Computational resources used for ColabFold predictions were provided by the e-INFRA CZ project (ID:90254), supported by the Ministry of Education, Youth and Sports of the Czech Republic. Identification of phosphopeptides from *in vitro* kinase assay was performed at the Turku Proteomics Facility supported by Biocenter Finland. We thank Dr. Julia Krasensky-Wrzaczek for her research input and constructive feedback on the manuscript.

## Author contributions

SJ and MW conceived the study and designed the research. AZ and SJ performed *in vitro* kinase assays and callose staining, imaging and quantification. AB, SJ and IK conducted FRET-FLIM experiment. SCS, AH and HN conducted and analyzed phosphoproteomics experiment. FC performed crossing. SJ and SL carried out physiological assays. MN and RP performed structural analysis. MP and JM carried out LC-MS/MS analyses. SJ conducted all other experiments. SJ wrote the first draft and MW edited and reviewed the paper. MW acquired the research grant. All authors participated in this research and contributed to the final draft of the paper.

