## Supplementary Figures for "CRK2 controls the spatiotemporal distribution of QSK1 at plasma membrane during osmotic stress"

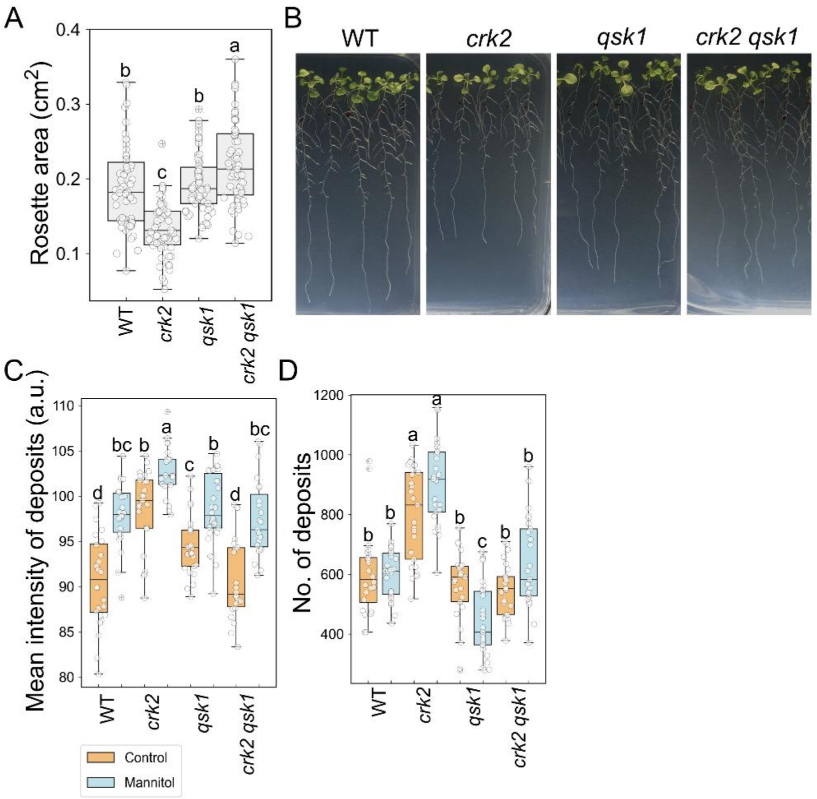


**Figure S1 (related to Figure 1). CRK2 and QSK1 regulate plant growth and osmotic stress response.**

(A) Rosette area of 12-days-old plants. (B) Root phenotype of the mutants. (C-D) Quantification of callose deposition in epidermal cells of aniline blue-stained leaves of plants treated with or without mannitol. The boxes represent the interquartile range (25th–75th percentile), the horizontal line indicates the median, the whiskers extend to 1.5× the IQR, and dots represent individual data points. Panel A, n=59, one-way ANOVA, Tukey's test; Panel C-D, n $\geq$ 22, two-way ANOVA, Tukey's multiple comparison test. See also Table S1 for panel A and Table S2 for panels C-D.

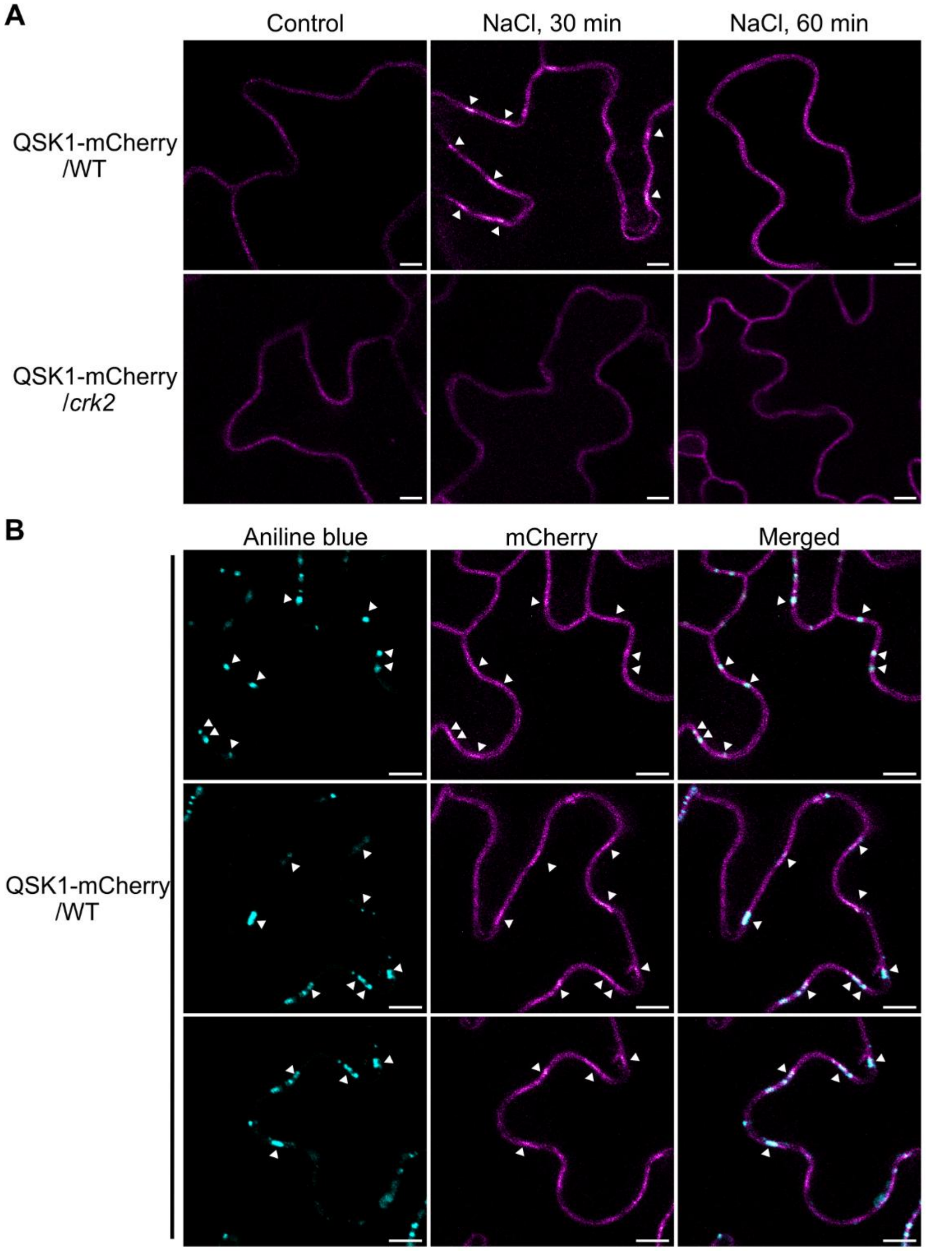


**Figure S2 (related to Figure 2). Enrichment of QSK1 at plasmodesmata requires CRK2.**

(A) Relocalization QSK1-mCherry in Arabidopsis leaf epidermal cells. Detached leaves from plants expressing QSK1-mCherry in WT and *crk2* backgrounds (QSK1-mCherry/WT, QSK1-mCherry/*crk2*) were treated with 150 mM NaCl and imaged at 30- and 60-minute post-treatment. Control samples were without salt treatment. In the QSK1-mCherry/WT plants, the punctate pattern of mCherry signal was prominent at 30 min but faded after 60 min of treatment. In the QSK1-mCherry/*crk2* plants, QSK1-mCherry did not enrich upon salt treatment. (B) Co-localization of aniline blue-stained callose deposits with enriched QSK1-mCherry signal showing that QSK1 enriched at PD upon osmotic stress treatment. Leaves were co-treated with 150 mM NaCl and 0.2 % aniline blue and abaxial surface was imaged with laser-scanning confocal microscope after 30 min of treatment. Objective lens 40.0 X oil immersion, emission wavelength for DAPI filter 461 nm and laser wavelength 405 nm. Detection wavelength 430-530 nm, laser transmittivity 2.0%, Scale bar 5 µm. Other laser settings for mCherry as Fig. 2.

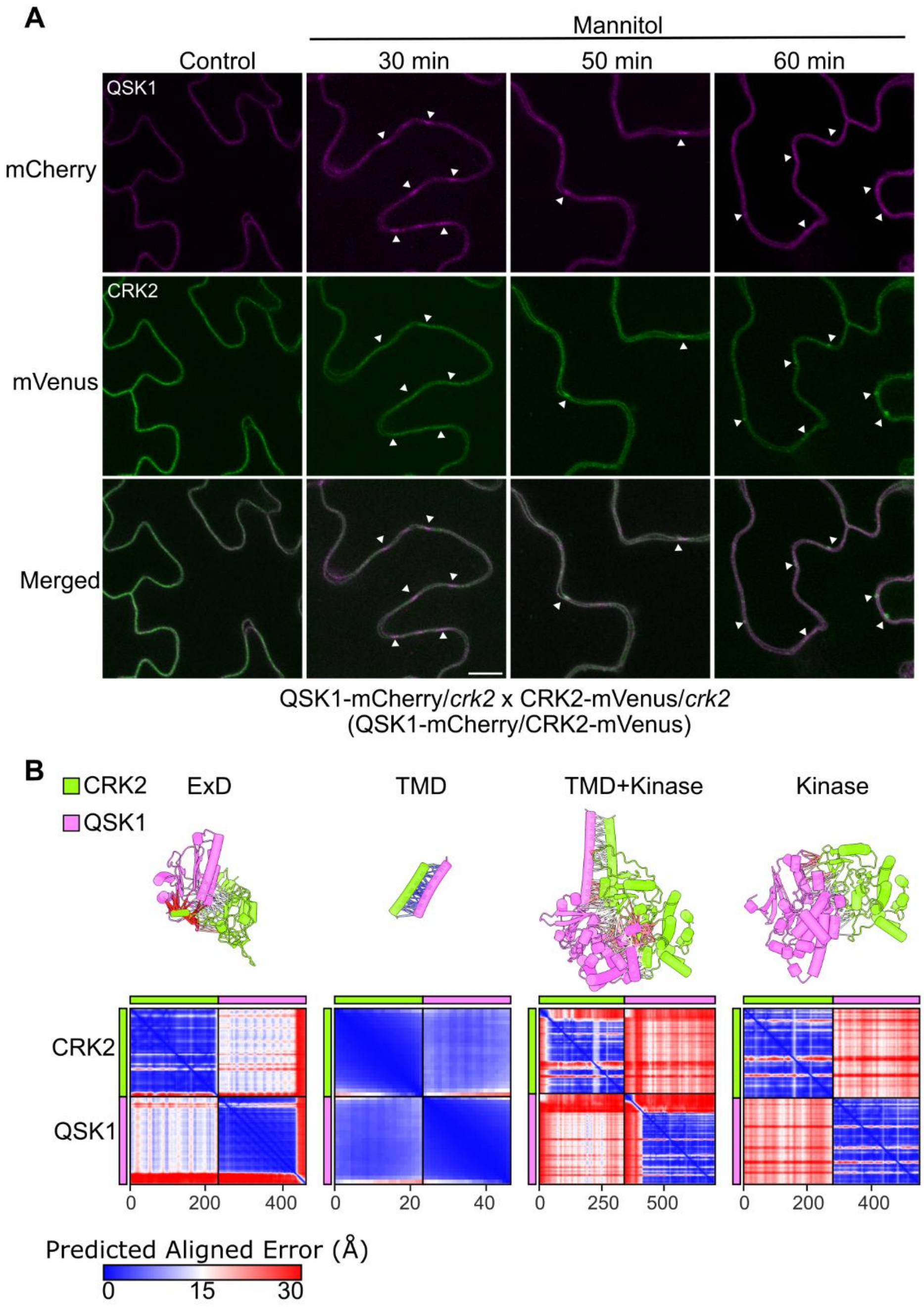


**Figure S3 (related to Figure 3). Spatio-temporal dynamics of QSK1 and CRK2 association at plasma membrane and structural modeling prediction of their interaction.**

(A) Arabidopsis leaf epidermal cells showing relocalization of QSK1 and CRK2 in the QSK1-mCherry/CRK2-mVenus plants. The images shown are captured within a window of 0-60 min of 0.4 M mannitol treatment. QSK1 enrichment is followed by CRK2 enrichment. Objective lens 40.0 X oil immersion. Laser wavelengths were 561 nm and 514 nm for mCherry and mVenus, respectively. Emission wavelengths were 610 nm and 527 nm for mCherry and mVenus, respectively. Detection wavelengths were 570-620 nm and 530-630 nm for mCherry and mVenus, respectively. Scanning of signals were done in a sequential manner to avoid signal cross-contamination between the channels. Scale bar 10 µm. (B) AlphaFold2 prediction of interaction between CRK2 and QSK1 segments- extracellular domains (ExD), transmembrane domains (TMD), TMD + Kinase domains, and Kinase domains. Predicted Aligned Error (PAE) plots show model's confidence. Lower PAE values indicate higher confidence of interaction. Lines are colored according to the PAE values. The modeled boundaries and predicated scores in Table S5.

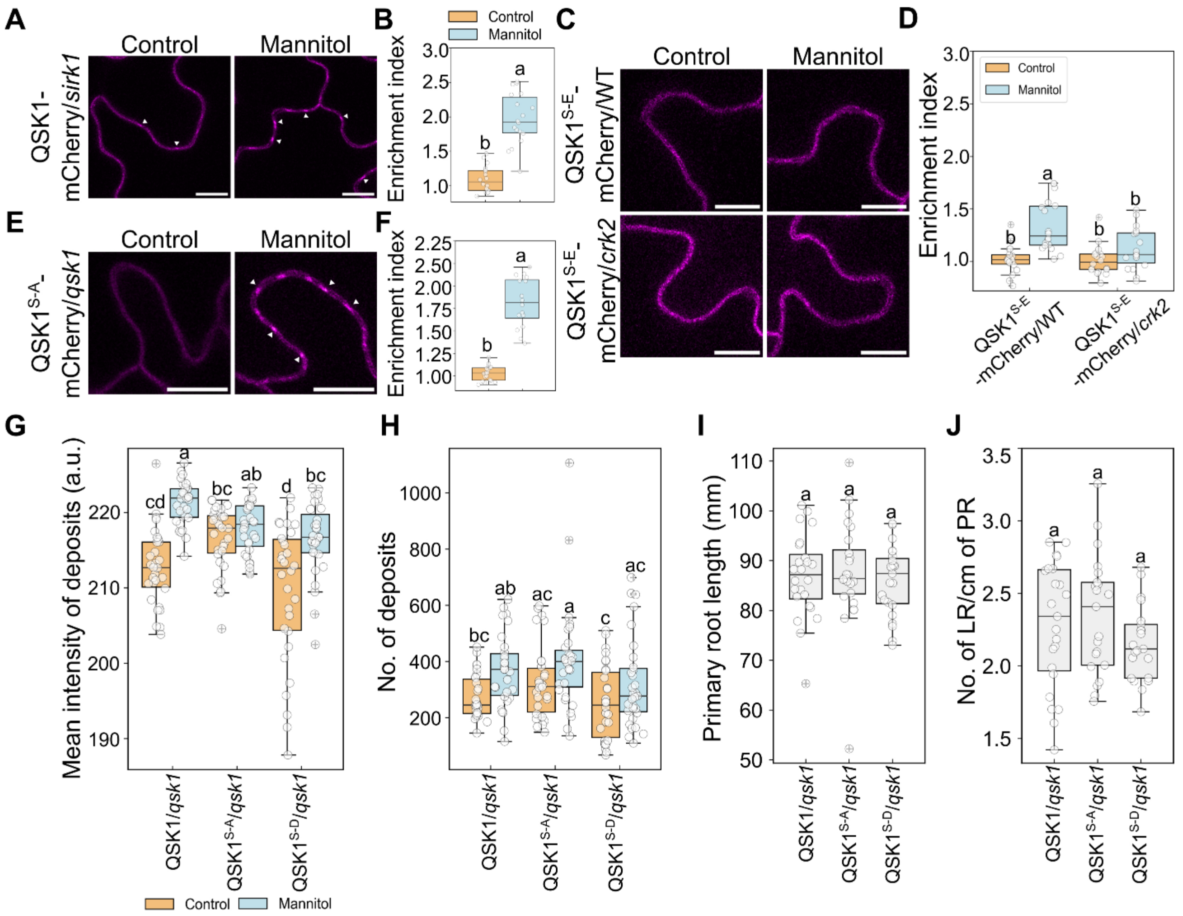


**Figure S4 (related to Figure 5). Phosphorylation of QSK1 at Ser-621 and Ser-626 inhibits its recruitment to PD and alters plant response to osmotic stress.**

Relocalization of QSK1-mCherry and QSK1 phospho-mutants in different genetic backgrounds of Arabidopsis leaf epidermal cells. (A-B) Enrichment of QSK1 at PD is independent of Sucrose-Induced Receptor Kinase 1 (SIRK1). Arabidopsis plants expressing QSK1-mCherry in *sirk1* mutant background show normal recruitment of QSK1 at PD upon mannitol treatment. A basal presence of QSK1 at PD was visible in control samples as well. (C-D) The phospho-mimetic QSK1$^{S-E}$ (QSK1$^{S621E-S626E}$) showed a weak enrichment at PD in WT background upon mannitol treatment. (E-F) Similar to WT background in Fig. 5C-D, the phospho-negative QSK1$^{S-A}$ (QSK1$^{(S621A-S626A)}$) showed high enrichment in the *qsk1* background upon mannitol treatment. In panels A, C- laser transmissivity 7.0%, objective lens 60.0 X oil immersion; panel D- laser transmissivity 2.0%, objective lens 40.0 X oil immersion. Scale bar 10 um. 3 independent transgenic lines were tested. Panel B, n=18, unpaired test with Welch's correction; Panel D, n=18, two-way ANOVA, Tukey's test; Panel F, n=17-18, unpaired test with Welch's correction. (G-H) Quantification of callose deposition in epidermal cells of aniline blue-stained leaves of plants treated with or without mannitol. $n \geq 22$, two-way ANOVA, Tukey's multiple comparison test. (I-J) PR length and LR density of QSK1 phospho-mutants at serines 621 and 626. The phospho-mutation of QSK1 at serines 621 and 626 does not impact the root phenotype under normal growth conditions. Panels I-J, n=22, Kruskal-Wallis test, Dunn's multiple comparison test. QSK1, QSK1$^{S-A}$ and QSK1$^{S-D}$ are the plants expressing WT, QSK1$^{(S621A-S626A)}$ and QSK1$^{(S621D-S626D)}$ variant of QSK1, respectively, in the *qsk1* mutant background. See also Table S3 for panels A-F, Table S8 for G-H, and Table S9 for I-J. Boxplots as described in Fig. 1 and Fig. S1.
